# A global dopaminergic learning rate enables adaptive foraging across many options

**DOI:** 10.1101/2024.11.04.621923

**Authors:** Laura L. Grima, Yipei Guo, Lakshmi Narayan, Ann M. Hermundstad, Joshua T. Dudman

**Affiliations:** Janelia Research Campus, Howard Hughes Medical Institute, Ashburn VA; Institute of High Performance Computing, A*STAR, Singapore

## Abstract

In natural environments, animals must efficiently allocate their choices across multiple concurrently available resources when foraging, a complex decision-making process not fully captured by existing models. To understand how rodents learn to navigate this challenge we developed a novel paradigm in which untrained, water-restricted mice were free to sample from six options rewarded at a range of deterministic intervals and positioned around the walls of a large (∼2m) arena. Mice exhibited rapid learning, matching their choices to integrated reward ratios across six options within the first session. A reinforcement learning model with separate states for staying or leaving an option and a dynamic, global learning rate was able to accurately reproduce mouse learning and decision-making. Fiber photometry recordings revealed that dopamine in the nucleus accumbens core (NAcC), but not dorsomedial striatum (DMS), more closely reflected the global learning rate than local error-based updating. Altogether, our results provide insight into the neural substrate of a learning algorithm that allows mice to rapidly exploit multiple options when foraging in large spatial environments.

## Introduction

Survival in natural environments depends on effective, adaptive decision-making across many simultaneously available resources - a process generally referred to as foraging. Foundational ethological models of foraging emphasize the relative quality of resources and their distribution in space as critical factors that shape foraging decisions [1]. Most notably, Charnov’s formalization of the problem - the marginal value theorem (MVT) [2] - proffers the fundamental insight that, due to the necessarily patchy spatial distribution of resources, foraging decisions can be framed as the choice of whether to ‘stay or leave’ an encountered resource. However, although many studies across species in behavioral ecology have found approximate agreement with the predictions of MVT [3–5], such a model makes a number of core assumptions, particularly with regards to animals’ knowledge of the environment [6]. Perhaps for these reasons, despite theoretical extensions to the standard form of the model [7,8], in the laboratory it has generally been necessary to train animals extensively before observing behavior consistent with MVT [6,9,10].

Other work in decision-making more broadly has focused on the dilemma of how to select between known options (more recently also referred to as ‘foraging’ [11,12]). Reinforcement learning (RL) models provide a compelling class of computational solutions able to address how agents make decisions when faced with multiple alternatives [13], for example by maintaining a running estimate of the quality of each option [14,15]. Such models not only provide an account of learning, but unlike MVT, RL is well-equipped to generate predictions on a choice by choice basis and has therefore been successful in relating observations of behavior and neural activity at the level of individual decisions. One particularly relevant example relates the phasic activity of midbrain dopamine neurons (mDANs) to the reward prediction error (RPE) term in temporal difference RL [16,17], connecting computations that determine how local option quality estimates are updated to animal choices as they evolve with learning [18].

However, as with MVT, RL as a model of decision-making has primarily been applied to the behavior of well-trained animals choosing between known, previously experienced options. Similarly, although RL formulations of decision-making can be extended to arbitrarily many possible options, studies have primarily focused on binary choice paradigms under an (often implicit) assumption that decision-making algorithms useful in this context can be generalized to more complex environments [14,19]. Thus probing decision-making in a many (>2) option context, beyond a greater ability to distinguish between models [20] and to leverage the paradoxically higher learning rates that animals exhibit in more complex environments [21], also offers the opportunity to explore what aspects of RL and foraging models are necessary for providing a good account of how untrained animals learn during naturalistic decision-making. Crucially, studying decision-making across many options also permits contrasting independent updating of option quality with variables applied across all options: in other words, is there a global parameter necessary for such learning? There is some evidence that striatal dopamine may play such a role in the updating of spatial value [22] but this has not yet been explored in the context of untrained animals foraging amongst many options over extended physical distances.

To address these questions we exposed untrained, freely-moving mice to six rewarding options of differing quality distributed across a large spatial environment while a moving gantry system allowed simultaneous fluorescence photometry. We began by first characterizing the emergence of structured behavior. We found that mice rapidly diverged from random decision-making and settled into a ‘matching’ strategy that, by taking into account the spatial configuration of options, resulted in near optimal rates of return in a majority of animals. We then constructed a unified mechanistic RL model that combines a ‘stay’ and ‘leave’ decision as in foraging theory with rapid learning of option qualities from experienced rewards. In this multi-option environment we found that a global, dynamic learning rate across options was necessary for model performance to match animal behavior. Finally, concurrent dopamine recordings using fiber photometry revealed that reward responses in the NAcC but not DMS were uniquely consistent with an implementation of this global learning rate term, both within a single session and with across day manipulations of option quality.

## Results

### Naive mice rapidly diverge from random behavior in a novel large multi-option environment

Completely untrained, naive, thirsty mice (total n = 23) were placed in a large, semi-dark arena (Fig. 1a, 1b; Supp. Fig. 1) of dimensions 213 x 45cm and challenged to discover, learn about, and exploit six options of differing quality in three-hour long sessions. The options were presented in the form of lick spouts arranged around the walls of the arena (Fig. 1b, right) which were replenished at deterministic, uncued intervals ranging from 30s at best to 2400s at worst (Fig. 1d) in two possible configurations experienced by separate groups of mice (Fig. 1c). The mice were free to select and travel between these options to check whether reward (4ul water) was currently available by licking at the spout. If reward was available, it was immediately dispensed. Otherwise, no reward was delivered. Crucially, rewards did not accumulate; any rewards not collected within the given interval were forfeited resulting in an opportunity cost to decisions made. Thus mice were presented with the challenge of learning how to effectively <u>Fo</u>rage across <u>M</u>ultiple <u>O</u>ptions: FoMO. To probe a potential neural correlate of learning, we applied concurrent fiber photometry to measure dopamine neuron terminal activity or release in two striatal subregions, the nucleus accumbens core (NAcC) and dorsomedial striatum (DMS) as animals engaged in FoMO (Fig. 1b, middle; Fig. 1d, bottom).

**FIGURE 1.**
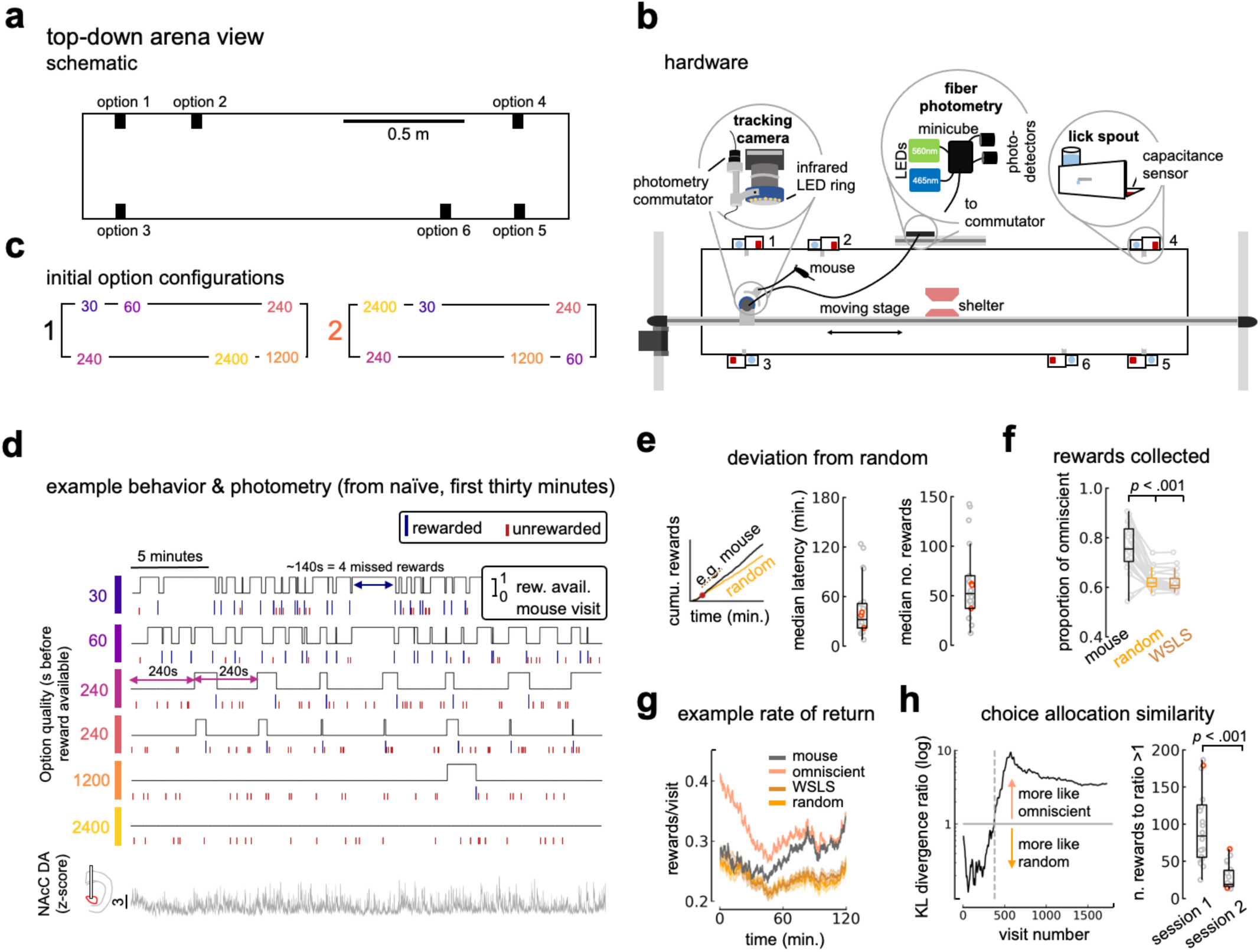
Untrained mice rapidly learn across six options in FoMO. **A** Top-down, to scale view of the arena with 6 options available from which to sample. **B** Top down illustration of key hardware in the arena. Mouse is to scale. **C** The two initial option configurations. Separate groups of mice experienced either config. 1 (left) or config. 2 (right). **D** Example mouse behaviour in the first thirty minutes of the first session, alongside real-time reward availability at each option. Blue long ticks and red short ticks indicate rewarded and unrewarded visits respectively. Bottom: concurrent fibre photometry recording in the NAcC. Note: all example data in this figure are from the same mouse. *Continued on following page.* **E** Left: example of mouse and associated random cumulative reward curves for a single run of the random policy simulation, with orange dot indicating calculated time that the mouse deviated from random. Center: median latency to diverge from random cumulative rewards curve. Right: median number of rewards to diverge from random policy in cumulative rewards. For both metrics, median is calculated over 10 runs of the random policy simulation. **F** Proportion of omniscient policy rewards collected for mice in comparison to random and WSLS simulations. **G** Rate of return (rewards/visits) over time for an example mouse in comparison to omniscient, WSLS, and random policy simulations. **H** Left: ratio of KL divergences for random to ideal for an example mouse over the space of visits. Dotted grey line indicates identified threshold for when animal choice behaviour is classified as more like omniscient than random. Right: number of rewards taken for animals to look more like omniscient policy simulation for sessions 1 and 2, median over ten simulation runs. Box plots represent median at center bounded by 25th and 75th percentiles of the data, with whiskers extending to the extent of the distribution barring outliers. All dots represent individual mice. Orange dots in E and H (right) indicate config. 2 mice. Significance testing: non-parametric Wilcoxon rank sum (h) or one-way RM ANOVA (f).

In contrast to trial-based tasks, FoMO allows for mice to freely sample each of the options available including returning to the same option without sampling from other spouts. We therefore distinguished consumptive licking bouts from the initial decision to visit an option by identifying a threshold over which the interval between consecutive licks was at least that of the minimum time to transition to a new spout, reflecting a putative new decision to resample the same option (Supp. Fig. 2). Segmenting behavior in this way yielded a stream of unique rewarded and unrewarded decisions to visit each of the 6 spouts (Fig. 1d). Similarly, mice were free to take indirect paths between options. We therefore developed a closed-loop gantry-mounted tracking camera system to record animals’ trajectories and allow for deeper characterization of behavior in FoMO (Fig. 1b, top left). This revealed that despite the size of the space available to them, mice typically traversed the full extent of the arena within minutes of exposure (Supp. Fig. 3a).

In session 1, as mice were completely naive, spouts were primed with a small drop of water to enable rapid discovery of all 6 options (Supp. Fig. 3b, left). Mice clearly differentiated between rewarded and unrewarded visits (Supp. Fig. 3b, right) and by the end of the first session had learned to quickly transition between spouts, averaging ∼800 (+/- 360 s.d.) visits across all options and resulting in a mean total of ∼350 (+/- 105 s.d) rewards (Supp. Fig. 3c). With additional experience on the same configuration in a second session, this increased to an average of ∼430 (+/- 58) rewards.

Even within the first sessions’ worth - 180 minutes - of experience animals exhibited dramatic changes in how they made decisions across options, indicating rapid learning. To benchmark animals’ performance we compared their choices over time to simulations of three simple decision policies: random choice as lower bound on performance, omniscient as upper bound, and win-stay lose-shift (WSLS) as a simple heuristic that can be effective in binary choice tasks. All simulations were constrained to the empirical choice times of individual mice in order to dissociate contributions of the decision policies *per se* from rate of sampling. Animals’ cumulative reward collection rapidly diverged from random performance within the first session - in most cases within 60 minutes (∼50 rewarded visits) (Fig. 1e). Consistent with learning from experienced rewards, the latency for individual mice to perform better than a random policy was correlated with the time taken to discover all 6 spouts (*spearman’s ρ:* 0.62, *p* < .004). In this context with six discrete options, WSLS did not perform substantially better than random. Thus mice outperformed both random and WSLS agents in their ability to collect available rewards across options (Fig. 1f).

Here the omniscient benchmark, as with the other simulations, did not have information about the timing of the reward intervals. Instead, it had access to the reward availability state of all six options at the time of the animal’s choices - and always took reward from one of the options if available - therefore acting as an upper limit to performance within the constraints of the individual mouse’s rate of behavior. Here we found, remarkably, that in some cases animals’ rate of return (rewards per visit) approached that of the omniscient simulation within the first session (Fig. 1g). Note that the initial decrease in rate of return across simulations and mouse data in this example is due to an increase in the number of visits over time at the beginning of the first session as animals became more familiar with the environment (Supp. Fig. 4, top).

This rapid increase in performance was presumably reflected by a change in how similar animals’ choices were to that of the simulations. To quantify this, we computed the KL divergence between mouse behavior and either the random or omniscient simulations, and then took the ratio between these metrics. A ratio over 1 indicates that animals’ sampling probabilities across options more closely resembled the omniscient benchmark than that of the random policy (Fig. 1h, left). The majority of mice distributed their choices in a manner more consistent with the omniscient simulation than random within the first session, and those that did not achieve this within the first 180 minutes did so in the second training session (Fig. 1h, right).

### Mice match their choices to relative reward across six options

We have demonstrated that, in FoMO, untrained mice were able to achieve a high rate of return with little experience: ∼30 minutes and ∼50 received rewards. However, although animal performance moved towards that of the omniscient upper bound, this analysis does not speak to how exactly animals achieved such performance. One possibility is that mice were attempting to increase the precision of visit timing at each option in order to collect each reward as soon as it was available. Indeed, there is evidence that animals can time their average behavior to learned intervals [23,24], albeit with extensive experience and a relatively high degree of error. However, this approach is unlikely here for two reasons. Firstly, the animals’ experienced reward rate at each port was much more variable than the true rate, with a large overlap of reward timing distributions across ports (Supp. Fig. 5a). Secondly, their visit timings did not correspond closely to the deterministic intervals of the underlying schedules (Supp. Fig. 5b).

On the other hand, the frequency with which animals sampled the individual options on offer changed rapidly with relatively little experience. After an initial period of random-like sampling, many mice began to distribute their visits in rank order of the reward schedules associated with each option (Fig. 2a). This allocation is reminiscent of a well-characterized behavioral principle known as ‘matching’, whereby organisms - from pigeons, to rodents, to humans - have been shown to choose options in proportion to how relatively rewarding they have been [25]. Indeed, during the first session, mice began to allocate their choices in a manner suggestive of matching (Fig. 2b-c). However, animals were not perfect matchers; they tended to ‘undermatch’ (Fig. 2b right, inset), a common cross-species observation [26–28]. In the current context, one driver of undermatching was animals’ relative naivety; as animals gained more experience, they moved closer to perfect matching both within (Fig. 2c) and across (Fig. 2d) sessions, as reflected by a change in the sensitivity of their choices to received reward [29]. Therefore, as far as we are aware, this is the first example of mice learning to approach perfect matching across six options within one or two sessions’ experience.

**FIGURE 2.**
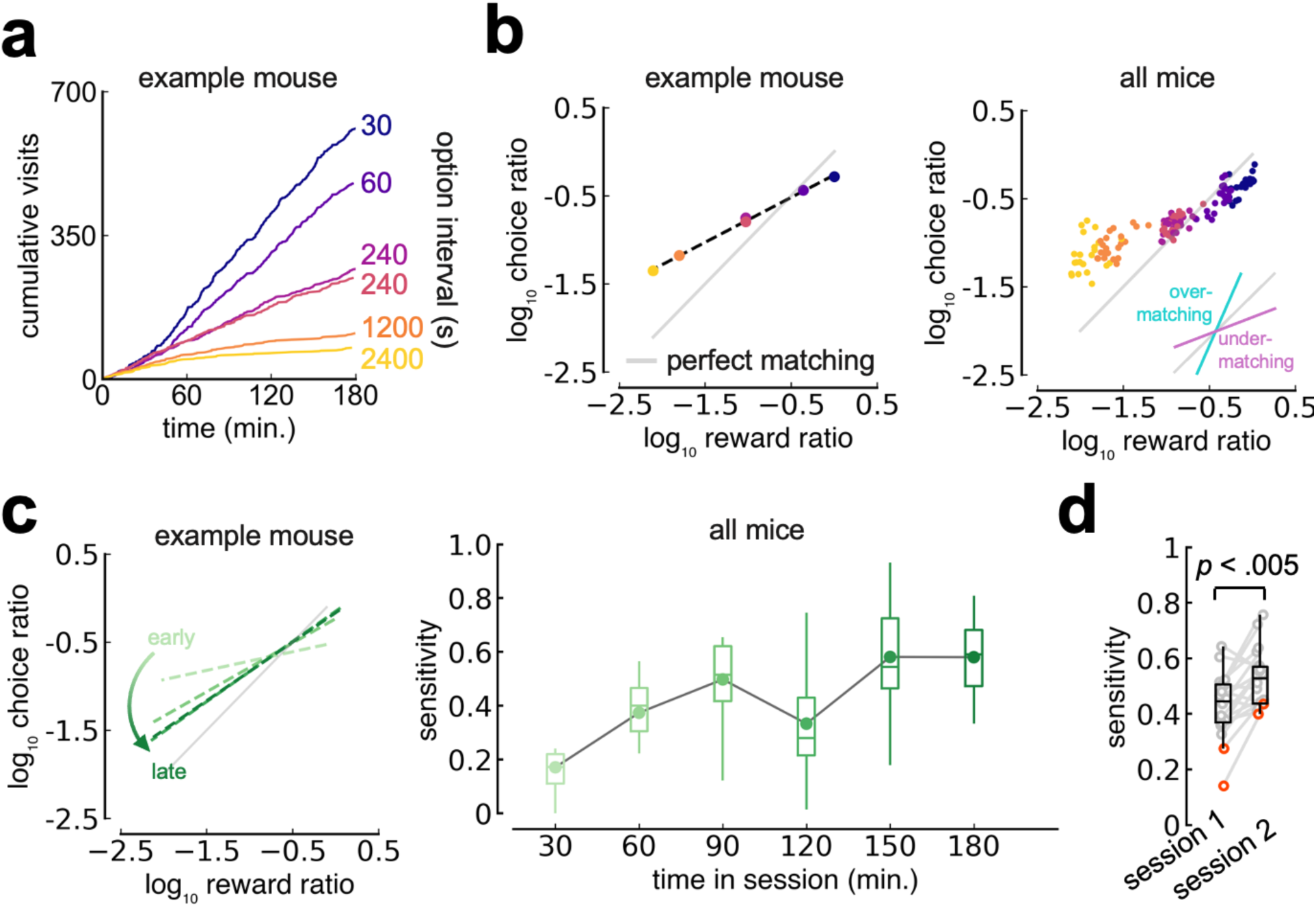
Mice learn to match their choices to reward across six options. **A** Example mouse cumulative visit distribution across 6 options. **B** Left, same mouse, one point per option reflecting choice ratio to reward ratio. Grey line indicates perfectly matching choice ratio to reward ratio. Right, the same but across the entire cohort, 6 points per mouse. Inset: example of under-matching vs. over-matching trends. **C** Left, example mouse (same as B) split into quartiles to show learning. Right, sensitivity across all mice in non-overlapping 30 minute moving windows. **D** Sensitivity slope comparisons for whole session 1 vs. whole session 2. Orange dots indicate mice that experienced config. 2. Significance testing: non-parametric Wilcoxon rank sum.

### Within a single session’s experience, mouse behavior reflects features of an optimal policy

Another unique aspect of FoMO is that, as in naturalistic foraging, mice have to select from options that are arranged in space. We conjectured that this might also shape decision-making and act as a possible driver of undermatching beyond amount of experience in the arena. Indeed, mice that were exposed to configuration 2 where the distance between the best options was greatest took longer to move towards perfect matching of choices to rewards (Fig. 2d, orange dots). With this in mind, we performed additional analyses to ascertain the degree to which physical distance between options, as well as absolute path length traveled, influenced decisions made in FoMO (Supp. Fig. 6). This revealed a number of insights pertinent to decision-making within a multi-option foraging context (Appendix 1), but in particular suggested that mice were especially sensitive to physical distance when choosing which option to sample from next (Fig. 3a).

**FIGURE 3.**
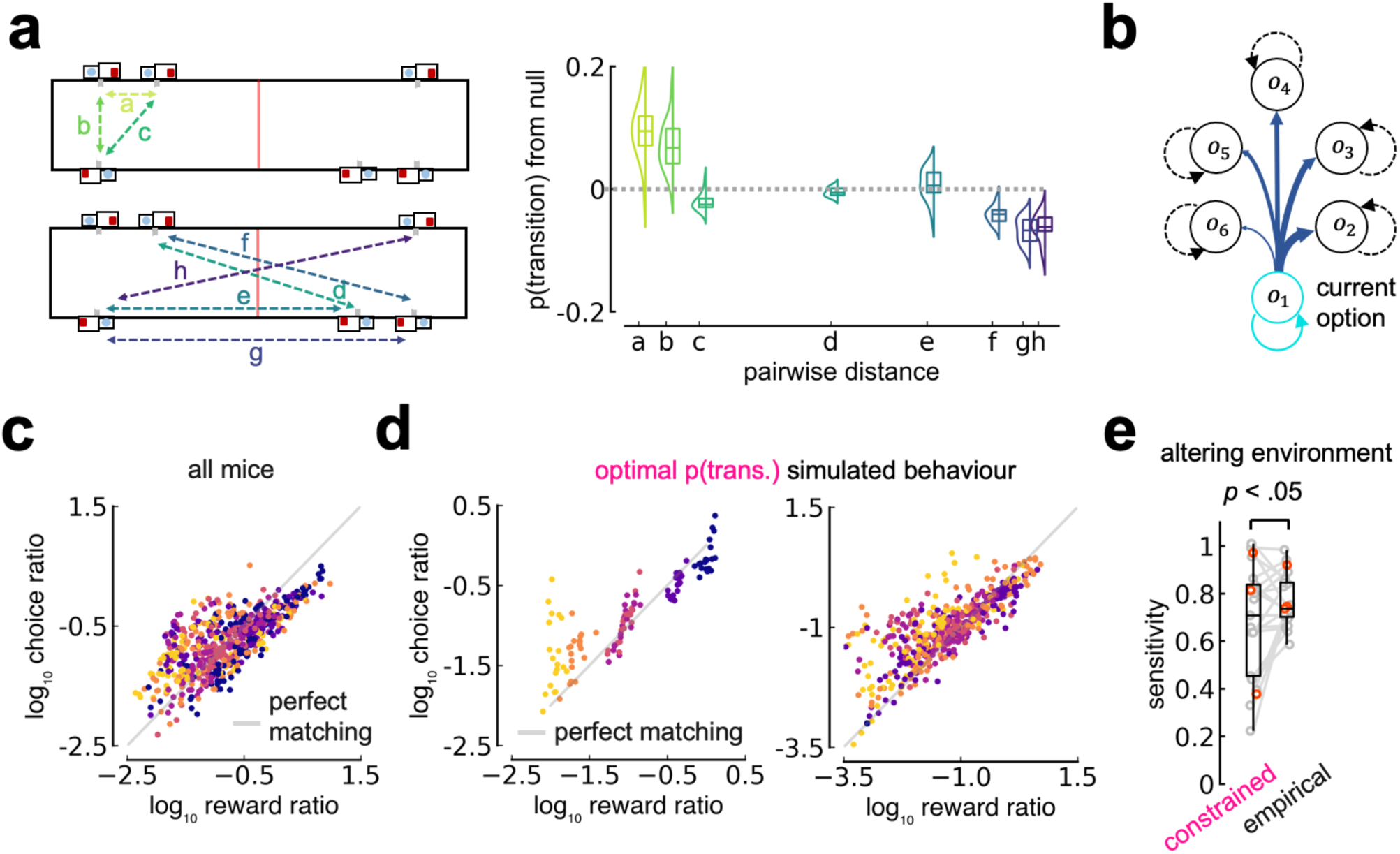
Mouse behavior reflects features of optimized transitions. **A** Left: all possible pairwise distances between ports, split by within the same half of the arena (top) and across the arena (bottom). Right: probability of making a transition by pairwise distance, relative to null distribution (overall probability of making the transition against all transitions). **B** Illustration of the set of transitions an animal or agent can make from each option, including returning to the same option. **C** Matching conditioned on current port, i.e. split by individual transition. Each dot represents one of 36 points for each animal. Colors represent port being transitioned to. **D** Agent behaviour generated by the optimal transition policy exhibits features of mouse behaviour, including undermatching (left) and conditional matching (right). **E** Matching sensitivity of agent behaviour when under environmental constraint or when sampling at the same timepoints as mice (empirical). All data plots are for full behavior from session 1. Significance testing: non-parametric Wilcoxon rank sum.

This sensitivity to distance suggests that animals might have treated the decision space in FoMO not as a vector of 6 possible options, but as a set of 36 possible transitions (Fig. 3b). Indeed, taking into account this transition structure by visualizing matching as conditioned on the current port - ‘conditional matching’ - results in behavior that more closely resembles perfect matching (Fig. 3c). This suggests that undermatching might in fact be a necessary consequence of behavior that aims to maximize reward when choosing between spatially distributed options. We tested this directly by first identifying the set of 36 transition probabilities that maximizes the reward rate, i.e. one form of an optimal policy, by way of a parameter search (see *Methods*). We then used this fixed policy as part of a generative model to simulate choice behavior. Crucially, the model was also constrained by physical distance between options. This was implemented by specifying the timing of the model’s visits to the empirical distribution of actual transit times for the specific transition being made. We found that the choice behavior of this optimality model exhibited both undermatching (Fig. 3d, left) and conditional matching (Fig. 3d, right), similar to the mice. Further, removing the constraint of distance between options resulted in model behavior closer to perfect matching (Fig. 3e). Therefore, undermatching in FoMO is not a flaw of animals’ approach to the task, but rather can be normatively accounted for as an integral feature of an optimal policy to maximize reward when foraging across many options.

### A reinforcement learning model that separately updates staying versus leaving and utilizes a dynamic, global learning rate explains mouse behavior

Although we provide a normative account for the observation of near ubiquitous matching behavior that has been observed across a range of species [30], the matching law has long been considered a ‘macroscopic’ description of behavior as opposed to a ‘microscopic’, or mechanistic, model of how decisions are made (*e.g.* [31]). In addition, the previously described optimality model applied a fixed, unchanging policy and so cannot speak to animals’ learning. Thus, we next sought to develop a mechanistic reinforcement learning (RL) model sufficient to account for the rapid learning of approximately optimal choice probabilities in untrained mice exposed to the FoMO paradigm.

One core aspect of RL models requires the agent to arbitrate amongst possible actions. In formal models of foraging, such actions include distinguishing between a decision to stay at a current option from pursuing a new option elsewhere [2]. While such an abstraction is not always used in models of decision making [32,33], here animals’ probability of staying at the best option was markedly lower than the probability of transitioning from the second best to the best option (Fig. 4b, left). This is reflective of an understanding of the open-loop, fixed interval nature of the schedules used. Our RL model therefore represented the decision to stay (P_stay_) separately from the decision to which alternative to transition (P_trans_; Fig. 4a-b).

**FIGURE 4.**
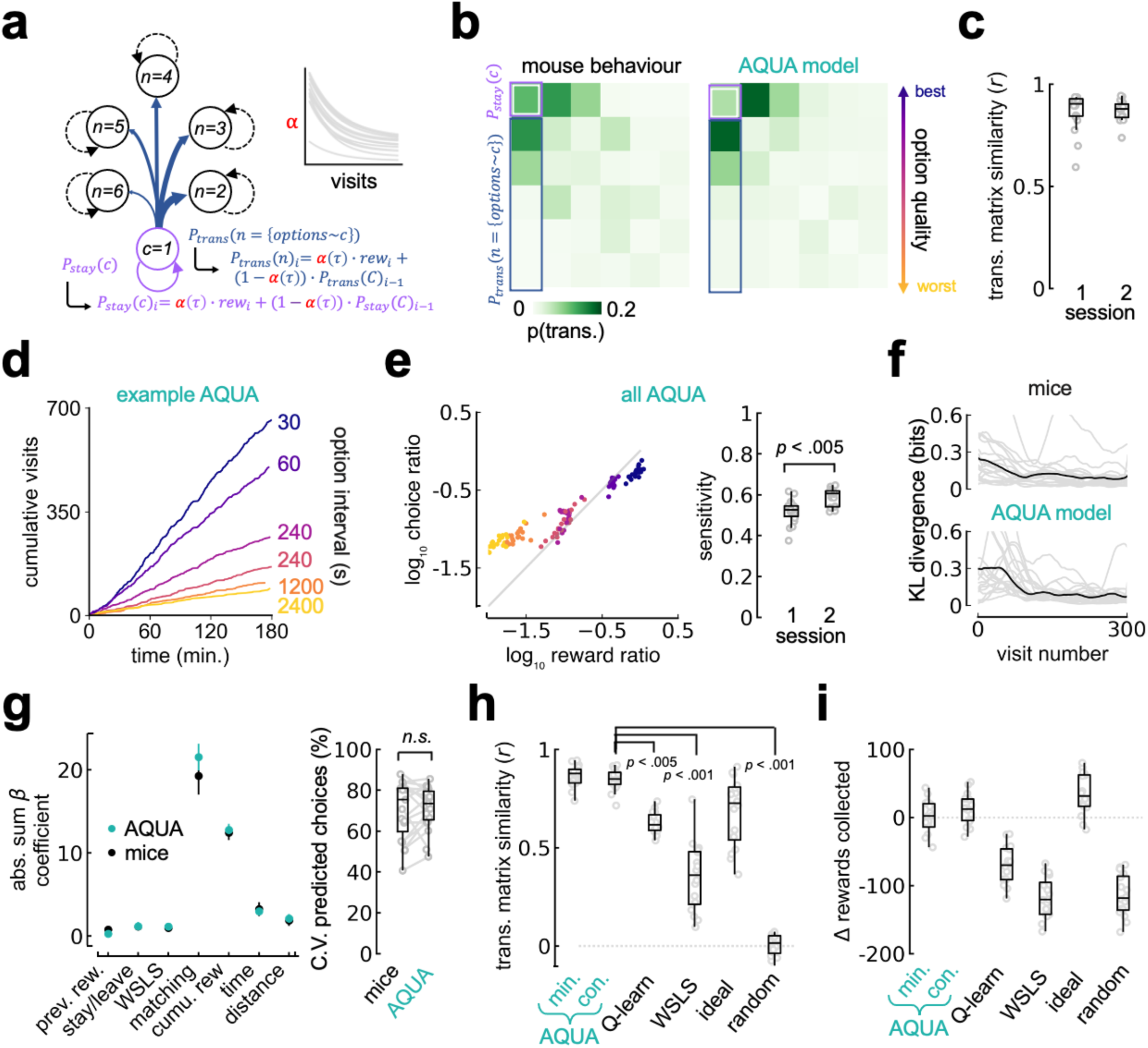
The AQUA model recapitulates mouse behavior with high precision. **A** AQUA model schematic. Inset: fitted optimal alpha across visits for each mouse. **B** Example transition matrices for a mouse and associated AQUA model run. **C** Transition matrix similarity between AQUA and mouse behaviour across sessions. **D** Example AQUA simulated cumulative visits. **E** Left: AQUA simulated matching curve across animals. Right: change in matching sensitivity across sessions with learning. **F** Change in KL divergence relative to a baseline of last 25% of decisions in the first session, for mouse (top) and AQUA model (bottom) behaviour. **G** Left: Full session 1 GLM comparing mouse (black) and AQUA (seafoam green) beta coefficients for the following predictors: whether option was previously rewarded; whether it was repeated (stay); win-stay lose-shift (interaction between prev. rew. and stay/leave); matching ratio value; cumulative reward; time since last visit to option; distance between prior option and current. Right: percentage of either mouse or AQUA choices correctly predicted by the GLM, with five-fold cross validation. **H** Transition matrix similarity across model comparisons. AQUA min. is best parameters found, AQUA com. takes the Center of Mass of the best 5% simulations. **I** Predicted rewards collected relative to animal behaviour. Significance testing: non-parametric Wilcoxon rank sum.

A second critical component of an RL model is that an agent must learn from received rewards and actions taken to estimate the quality of available options in the environment. The vast majority of RL models apply value learning, whereby a utility estimate is updated in proportion to the difference between the current estimate and reward - the reward prediction error (RPE) [13,17]. Of note is that as long as this update proportion (generally referred to as the ‘learning rate’, ***α***) is small, such models can produce matching behavior [34]. This comes at a cost, however, as models that use a constant, small value of ***α*** learn slower than observed behavior in animals [34,35], and here we found learning to be especially rapid. Alternatively, there is no requirement that ***α*** be static. For example, it is common practice to exploit adaptive learning rates when training models in machine learning [36,37] and there is increasing recent evidence for neuromodulatory control of learning rate in animal brains [14,38]. If we allow a dynamic learning rate, ***α***(***τ***) where ***τ*** is some measure of experience in the environment, it may be possible to develop a model that both rapidly learns an approximation from limited experience and recreates the matching behavior observed (Supp. Fig. 7a).

We thus simulated agents that learned an incrementally updated estimate of option quality with an optimally fit parametric ***α***(***τ***) function where ***α*** is comprised of both a static offset and dynamic exponential decay, either of which can span {0,1} (Fig. 4a; see *Methods*). Such agents perform a computation akin to a type of RL model known as Q-learning [13,39], but scaled by distance to convert “values” to utility and updated using an adaptive learning rate. To indicate the multiple insights that are combined into this model we refer to it as **A**daptive rate **Q**-learning of **U**tilities for **A**pportioning actions (or ‘AQUA’).

We first characterized the performance of AQUA agents with optimized parameters governing the time evolution of ***α***(***τ***) as evolving across visits. Such agents provided remarkably good accounts of behavior both across multiple timescales and in terms of its higher order structure. For example, AQUA models can account for a substantial majority of the variance in the 36 dimensional transition matrix across all ∼20 mice (Fig. 4b-c; session 2: r^2^=74.8 ± 8.2%). AQUA also produced quantitatively similar undermatching behavior both on a single session (Fig. 4d; Fig. 4e, left) and across session basis, with a sensitivity that increased over the first two sessions (Fig. 4e, right) in a manner remarkably similar to data (Fig. 2b-d). To probe whether AQUA captured the rapid change in behavior early in learning, we visualized the evolving probability of choosing each option relative to stable behavior in the last quarter of the first session by calculating the KL divergence over decisions. Mouse and model behaviour changed at a similar rate over time (Fig. 4f). Finally, we constructed a generalized linear model (GLM) using multinomial logistic regression to predict the probability of a given choice across six options on a decision by decision basis for both mice and the AQUA model given a set of relevant predictors (see *Methods*). AQUA produced a remarkably similar statistical structure to mouse behaviour (Fig. 4g, left) such that the cross-validated accuracy of the GLM was statistically indistinguishable when comparing datasets (Fig. 4g, right).

We next compared AQUA to several other models bounded by the baseline of a random agent that doesn’t learn and the ceiling of an ‘omniscient’ agent that always takes a rewarding option if one is available - the same approach used in the simulations in Fig. 1. To facilitate comparison we examined both the explained variance of the transition matrix (Fig. 4h) and the predicted number of rewards collected in a session compared to the observed reward collection count (Fig. 4i). For comparison to other proposed models which constitute sub-components of AQUA we also simulated a standard Q-learning model [14] and a WSLS strategy - both of which have been argued to explain behavior in binary foraging and decision-making tasks [19]. AQUA with a dynamic learning rate explained the most variance in the transition matrix and yielded the most accurate prediction of the number of rewards collected compared to all other simulations (Fig. 4h-i; Supp. Fig. 7b).

### NAcC dopamine reflects the AQUA dynamic learning rate

At this point we have described key features of animals’ behavior in FoMO, including rapid learning through adoption of a matching strategy that enables animals to achieve near optimal foraging performance. We can explain a large proportion of this behavior with a mechanistic RL-based model that combines key features including accumulating estimates of option quality, weighting by distance cost, and a global dynamic learning rate. To probe the physiological correlates of these behaviorally-inferred learning mechanisms, we recorded dopamine terminal activity (using GCaMP8m) or release (using dLight1.3b) in two regions of the striatum, the NAcC and DMS, during this behavior (Fig. 5a; Supp. Fig. 8). Signals recorded using either strategy looked similar in this context (Supp. Fig. 9), and so we collapse across both approaches for all analyses here.

**FIGURE 5.**
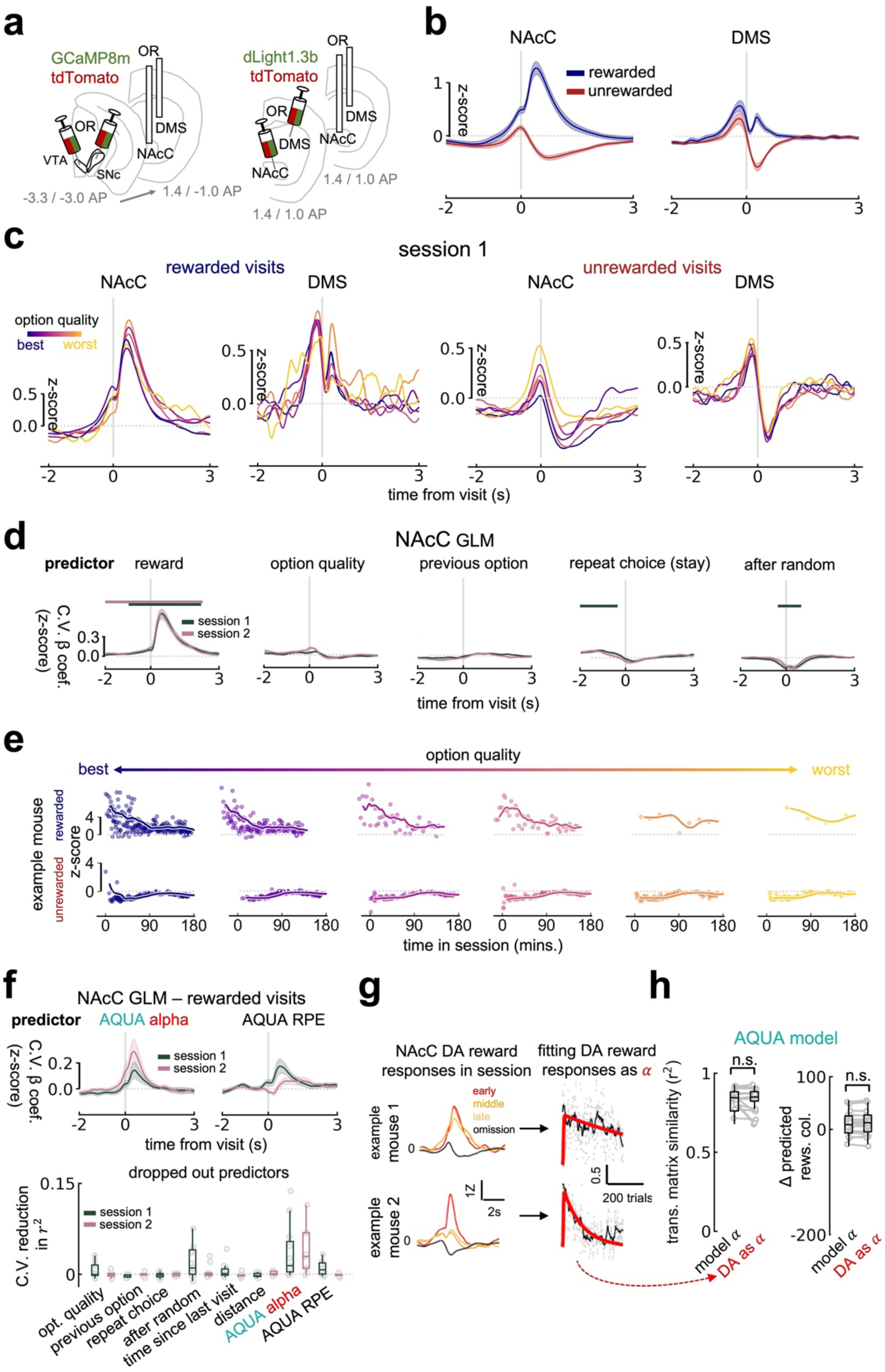
NAcC dopamine reflects the AQUA global learning rate. **A** Viral and fiber implantation strategies for fiber photometry recordings, using either a calcium sensor (GCaMP8m) or a dopamine sensor (dLight1.3b) in either the NAcC or DMS. **B** Mean z-score response to rewarded and unrewarded visits across regions across sessions 1 and 2. **C** Mean z-score response to rewarded and unrewarded visits across regions in session 1 split by option quality. Traces are baseline subtracted to the start of the plotted period for visualization. **D** Five-fold cross validated z-score predictor weightings (beta coefficients) over time for a selection of key predictors for a linear regression to predict NAcC dopamine in both rewarded and unrewarded visits on a sample to sample basis. Bars indicate significant difference from zero (*p* < .01), Benjamini-Hochberg corrected for multiple comparisons. **E** Example of all rewarded and unrewarded visit responses, split by option, across session 1. Points are peak response to reward or lowest point in dip in 0-2s period from visit. Line is interpolated rolling mean with 2000s window. **F** Top: Same as in *d*, but GLM is now run on rewarded visits only and includes AQUA-derived alpha and RPE. Bottom: 5-fold cross-validated reduction in GLM r2 with predictor drop out, predicting the mean of the peak response in the 1s period after reward delivery. AQUA alpha was the only predictor that resulted in a significant reduction in r2, *p* = .02, Benjamini-Hochberg corrected for multiple comparisons. **G** Illustration of use of NAcC DA reward responses, fit on a per mouse basis, to estimate AQUA alpha, for two example mice. **H** Comparison of model-fitted alpha performance with NAcC DA as alpha for transition matrix similarity to mice (left) and delta predicted rewards collected to mice (right). All photometry trace plots: lines are mean, shaded error bars are SEM. Significance testing: non-parametric Wilcoxon rank sum.

As shown by a wealth of previous literature, here we found that the signal in both the NAcC and DMS robustly responded to water delivery when a visit was rewarded, and dipped below baseline after an unrewarded visit (Fig. 5b). However, inconsistent with proposals that dopamine encodes the value of options within the context of decision-making [18], we did not initially find explicit rank order scaling of reward responses to the objective option schedules within the first session in either region (Fig. 5c, left), even when limiting to after animals had deviated from random (Supp. Fig. 10a, left). In fact, responses for the best and worst options were highly overlapping, despite the former representing up to a 80x higher rate of reward and several fold higher probability of reward per visit. This was also the case for session 2 reward responses, where animals were more stable in their decision-making behavior (Supp. Fig. 10b, left). Reward responses to each option were also not consistently reflected by the subjective estimate of option quality, experienced reward rate, in either session 1 or session 2 (Supp. Fig. 10c).

In contrast, unrewarded visits did show scaling in roughly rank order of option quality (Fig. 5c, right) particularly in session 2 (Supp. Fig. 10b, right). However, due to the nature of the task - as well as prior evidence that dopamine responses can correlate with a broad range of variables [40–42] - it is likely that responses at each visit here reflect a combination of variables that are also changing over time as mice learn. To identify the independent contribution of such variables to dopamine signals we ran a regression analysis to predict the dopamine moment by moment, and as it evolves across the session (see *Methods*). In a first model, responses to both rewarded and unrewarded visits were included. In the NAcC, as expected from 5b and 5c, reward but not option quality had a large positive influence on the dopamine response (Fig. 5d). Inspired by our behavioral findings, we also investigated predictors including the quality of the previous option, stay versus leave decisions, and learning phase (before or after the random-like period of sampling). Yet when multiple variables were considered, only the phase of animals’ learning was reflected in the phasic NAcC visit responses; responses after random were depressed, particularly in the first session (Fig. 5d, right). DMS regression weights were altered in response to each of these predictors in the same direction and with a similar magnitude as in the NAcC (Supp. Fig. 11a).

The correlation in dopamine response magnitude with learning phase - before vs. after random - prompted us to visualize rewarded and unrewarded visits across the session. This revealed that the magnitude of NAcC responses to rewarded visits in particular decayed with experience, and the rate of this decay was similar across all options (Fig. 5e). At face value this pattern is strikingly reminiscent of the change in ***α*** as fitted by the AQUA model to mouse behavior, initiating at a high value that relaxes over time (Fig. 4a, inset). Indeed, a similar pattern is apparent in NAcC dopamine responses in other datasets, albeit in rats with more training and sampling across fewer options (Supp. Fig. 12; [22]) This effect was also dissociable from changes in engagement across the session, which did not decay across the three hours (Supp. Fig. 4). In addition, a similar change in response magnitude was observed in unrewarded visit responses, with the largest pauses in NAcC dopamine observed early in the session and reducing in magnitude with experience (Fig. 5d). Equivalent formulations of AQUA update rules can utilize learning from unrewarded visits (see *Methods*). While we do not explore this in detail here, this is consistent with recent evidence that pauses in dopamine firing slow extinction learning [43].

We therefore included a dynamic learning rate derived from AQUA as a regressor into a reward-specific GLM. Usefully, the update rule used in AQUA can be formulated in algebraically-equivalent forms that either do or do not compute a reward prediction error (see *Methods*). We took advantage of this to also include AQUA-derived RPE values as an additional separate predictor of striatal dopamine signals that, in contrast to the global learning rate, are locally updated to each option. Dopamine in the NAcC (Fig. 5f, top) but not the DMS (Supp. Fig. 11b) was predicted to some extent by both AQUA ***α*** and RPE, but an additional stringent drop-out test revealed that ***α*** reduced cross-validated explained variance of the model more than RPE in both sessions 1 and 2 - and indeed was the only predictor to do so with statistical significance (Fig. 5f, bottom).

Intriguingly, removal of any of the predictors from the model had little effect on GLM performance in explaining variance of DMS signals (Supp. Fig. 11c). To exploit the unique context in which these recordings were made we also investigated the distribution of transients across the arena. Here DMS responses were consistently more distributed across space than the NAcC (Supp. Fig. 11d, left). This difference was confirmed by calculation of the entropy of response distributions (Supp. Fig. 11d, right). Although the bulk of our striatal dopamine analyses here focus on outcome responses, we highlight this feature as warranting future exploration.

As a final test of whether observed dopamine dynamics from the photometry recordings in NAcC could be consistent with the dynamic learning rate term ***α***(***τ***) from simulations of AQUA, we fit the magnitude of reward responses for each reward with the same parametric function used to describe ***α***(***τ***) and found that it could capture response dynamics over the course of the session (Fig. 5g, see *Methods*). Next, we used these inferred parameters of ***α***(***τ***) from dopamine responses (‘***α***==DA’) to re-run simulations and compare predicted behavior using dopamine responses to the optimal fit (‘***α*** min’). We found that assuming dopamine signaling instantiates ***α***(***τ***) in AQUA provides an equivalently good accounting of behavior to an optimal fit of ***α*** (Fig. 5h). Thus, our data are consistent with the conjecture that mesolimbic dopamine signaling implements a global adaptive learning rate.

### Mice, the learning model, and dopamine adapt with across day changes to the environment

Our model-based analysis suggests that mesolimbic dopamine responses to reward delivery are most consistent with an implementation of a learning rate that is initially large early in a session and slowly relaxes to a low value - a form of dynamic learning rate that is suitable for effectively apportioning decisions across many options. Although our previous analyses were able to dissociate error computations and an adaptive learning rate to an extent, it is still difficult to resolve these differences because error computations and dynamic learning rates converge significantly when option quality is static. We reasoned that introducing changes to option quality should make the predictions of these two functional models more distinct.

To examine this we exposed mice to up to three further sessions in which the reward schedule assigned to each option changed (Fig. 6a). Mice were able to rapidly adapt their decisions to changes in option quality in a way that maintained approximate matching of choice prevalence to integrated reward (Fig. 6b-c). The AQUA model tracked changes in structure of options with similar behavior to the mice, exploiting dynamic learning rate modulation in each session. To compare the evolution of behavior around a single transition, from session 2 to 3, we computed the difference in the probability of choosing each option relative to stable choice probabilities from the end of session 2 as KL divergence. Both mice and AQUA simulations’ change in choice probabilities peaked within the first ∼50 visits of session 3 (Fig. 6d).

**FIGURE 6.**
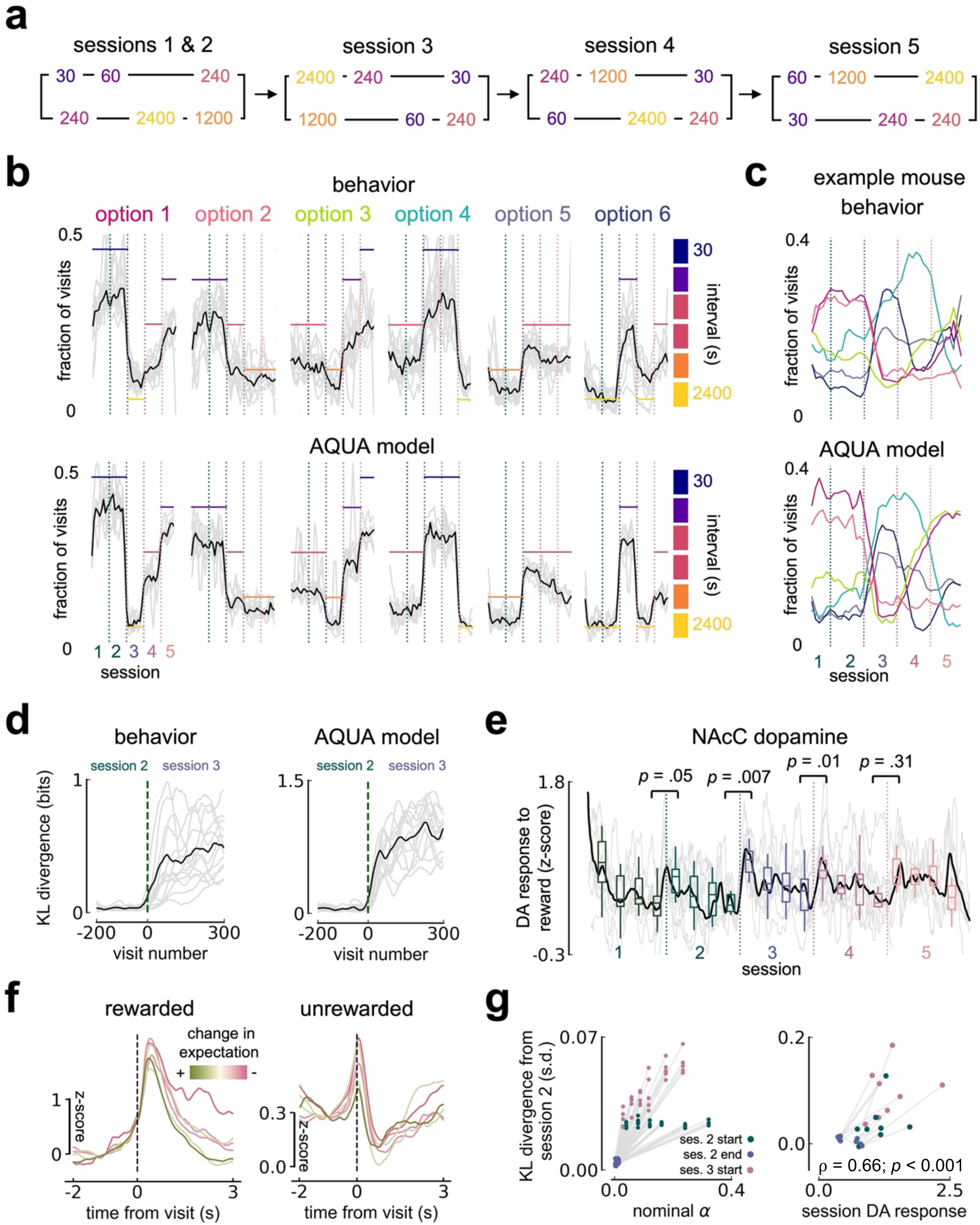
Mice, AQUA, and NAcC dopamine track schedule changes across days. **A** Change in assignment of option qualities to spouts across days. Note that only animals that experienced config. 1 underwent this additional manipulation. **B** Fraction of visits to each option across sessions for mouse behaviour (top) and AQUA model simulations (bottom). Black line is median across all animals. Grey lines represent individual mice. All lines smoothed with a running window of 20 minutes. Colored horizontal lines indicate option quality within the given session (right y axis). **C** Example mouse behaviour (top) and model simulation (bottom) for fraction of visits for each option across sessions, smoothed with a running window of 20 minutes. **D** KL divergence from session 2 to session 3, relative to average session 2 distribution of choices for mouse behaviour (top) and AQUA model (bottom). Black line is median and grey lines are individual mice, linearly interpolated and then smoothed with rolling window of 50 visits. **E** NAcC dopamine z-score response to reward collapsed across options, across sessions. Reward response is calculated as peak within 2s window after visit. Black line is median, grey lines are individual mice, linearly interpolated and smoothed with a rolling window of 500s. Overlaid boxplots represent quartiles with whiskers extending to the rest of the distribution. *P*-values calculated from Wilcoxon signed-rank test comparisons of quartile values. **F** Mean z-score response in NAcC to rewarded (left) or unrewarded (right) visits for each option in session 3 colored by change in expectation, i.e. if option quality in session 3 > option quality in session 2, the line is green. Averaged across all mice. **G** KL divergence comparisons from session 2 relative to nominal alpha (left) or mean session DA response (right), split by session 2 start or end, or session 3 start. Each point is one mouse.

The first time option qualities changed, session 3, we observed a pattern of mesolimbic dopamine responses to reward delivery that peaked early in the session and gradually decayed over time, similar to sessions 1 and 2 (Fig. 6e). The increase around the transition from the end of session 2 to the beginning of session 3 was uniquely reliable in all mice as compared with changes between session 1 and 2 where there was no change in option quality, or the transition between sessions 4 and 5 where there was a smaller overall change in quality (Fig. 6e). However, putative errors between expectation and reward availability were neither reflected in differential responses of mesolimbic dopamine activity to reward delivery, nor in differential decrease of activity following omitted rewards from previously high quality options (Fig. 6f). Thus dopamine dynamics reflected a global change in the landscape of options, but not specific mismatches between expectation and outcome at each port.

How a learning rate should relate to changes in the behavioral policy are distinct from errors. A high learning rate effectively makes option choices more sensitive to recent history of observed rewards. This can lead to differences in observed behavior even across periods where there is not a global change in option quality. To illustrate this point we can compare simulations of the AQUA model across a set of 3 epochs that should be rank ordered with respect to increasing changes in behavioral choice, relative to the end of session 2. Late in session 2 when learning rate is low and options are stable, choice behavior should change little. In contrast, even though options are stable early in session 2, learning rate is relatively higher at this point and should therefore produce greater KL divergence. Finally, early in session 3 a combination of both high learning rate and a global change in option qualities should make choice behavior most dynamic. We can confirm these expectations about model behavior by running a range of simulations across a broad set of scaled learning rates (Fig. 6g, left). If mesolimbic dopamine responses to reward reflect the setting of the learning rate, then we should expect that this same correlation between learning rate and choice variability exists when replacing known alpha with observed dopamine data. Indeed, there was a robust correlation between NAcC dopamine responses and observed behavioral choices for animals that experienced the session 2 to 3 transition (Fig. 6g, right).

## Discussion

Here we established a novel paradigm, FoMO, to understand how completely untrained and naive animals learn about and exploit many (6) concurrently available options distributed over relatively large spatial scales (∼2.1m). Mice rapidly learned to sample options in proportion to the history of received reward, referred to as ‘matching’ behavior, which achieves near-optimal rates of reward in FoMO. To gain mechanistic insight into this rapid learning, we developed a model that combines key insights from foraging theory and reinforcement learning; namely, a dynamic global learning rate, online updates of option quality weighted by travel costs, and an independent estimate of the utility of persisting in the current option. We show that across multiple metrics this model, AQUA, was able to recapitulate behavior of all individual mice with exceptionally high accuracy. This comparison suggests that the key components of AQUA may provide a good account of the underlying computations that guide mouse learning. Consistent with this conjecture, we found that the dynamics of dopamine responses in the core of the NAcC, but not the DMS, were highly similar to the global learning rate term in AQUA. Although other local accounts of dopamine signaling (*e.g.* prediction error) can be quantitatively similar to a dynamic learning rate in a stable environment, abrupt switches in option arrangement across sessions provided additional strong evidence that phasic NAcC dopamine reward responses are most consistent with a dynamic learning rate during foraging.

The learning seen in FoMO occurred with no prior pre-training or exposure to the environment. Yet animals quickly settled into near-optimal behavior remarkably quickly, at least an order of magnitude faster than previous studies that featured fewer options [14,22,44]. In addition, mechanistic accounts of how animals learn about the quality of alternative options available to them is an understudied aspect of foraging theory. Heuristic approaches, such as MVT, generally assume prior knowledge of key properties of the environment such as the average quality of all options or the time required to travel to new options (although attempts have been made to tackle this issue in optimal foraging, e.g. [45]). Here we show that dynamic changes in incremental updating of option quality into running estimates weighted by the distance between options is sufficient to balance a rapid emergence of structured behavior with stable, near optimal choice behavior with extended experience.

Due to its parsimony, MVT and variants thereof remain the most popular model to explain foraging decisions when studied in the lab. However, whilst it provides beneficial insights into computation in such contexts, MVT usually provides a qualitative fit of behavioral trends and speaks less to individual decisions and variance across animals [44]. In addition, the applicability of MVT is restricted to a specific form of environment that is not known to mimic natural statistics. Indeed, it is readily apparent that a decision heuristic based on MVT would generalize poorly to FoMO and other environments with patch statistics that are not monotonic, exponentially-depleting [46]. In contrast, AQUA not only explains initial learning and decision-making in FoMO, but because it incorporates a core principle of MVT - directly estimating the utility of persisting at an option - it also provides predictions consistent with MVT when exposed to depleting patch environments (Supp. Fig. 13a) and reproduces matching similar to that observed in two-armed bandit foraging (Supp. Fig. 13b). However, AQUA provides additional insights into distinct computations and putative neural representations of decision variables that could determine variable patch residence times and undermatching phenomena that warrant future study. This further demonstrates the power of combining core insights from reinforcement learning and foraging theory into a single model and suggests that relying on heuristic models tuned to specific assumptions about environment structure, whilst critical for developing insights initially, are more limited in their explanatory power. This highlights the need for continued conceptual development to better understand how diverse organisms forage in more complex environments.

Our work here is not the first attempt to express either MVT or Herrnstein’s matching law as mechanistic, agent based models using methods from reinforcement learning [44,47,48]. Previous work has argued that in some cases simple heuristics can provide a more efficacious and parsimonious account of foraging decisions than a model-free RL equivalent [47], although this depends upon certain assumptions regarding environment dynamics [49]. It has also been suggested that combining model-free and model-based components might be an approach both flexible and computationally simple enough to generate adaptive and effective foraging decisions [50]. Indeed, here we successfully took inspiration from a classic model-free form of RL, Q-learning, and combined it with additional model-based information such as distinguishing decisions between staying at the same option and leaving to go elsewhere. However although with AQUA we chose to represent each option with an independent estimate of quality - and have some evidence that mice do the same (e.g. 240s options are sampled equally even when they are grouped with different quality options) - future implementations of AQUA may benefit from a more explicit representation of hierarchical environment structure.

Foraging theory is defined both by a requirement that the organism make sequential decisions [33] - a core feature of the FoMO paradigm - but also by an appreciation that an agent must ‘travel’ between dispersed options in a patchy environment. The inferred cost of travel is a critical part of the decision to stay or leave a currently sampled option. An elegant insight in MVT is that if the quality of a current option is computed as a reward rate, then it can be compared to the cost of travel duration to find an optimal leaving time. However, it is unclear whether the cost of travel for an animal can be fully captured by duration while failing to account for path specific features such as distance, navigational difficulty, physical effort, risk of predation, etc. As far as we are aware, this assumption of travel time being uniquely critical has not been explicitly tested, and manipulations of travel time are rarely disambiguated from distances in practice [9]. The large spatial environment in FoMO was sufficient to observe travel times (∼1-20 sec) that cover the range of simulated travel times used in previous paradigms in the lab; however, travel time was only moderately correlated to distance between options. Although mice in FoMO were indeed sensitive to the spatial distance between options, they did not obviously attempt to minimize path lengths nor travel time *per se* and neither were critical predictors of behavior. This may partly reflect their relative inexperience navigating in the dark FoMO enclosure (a few hours) and/or competing desire to completely explore the arena (∼100cm^2^). In future work it may be critical to study foraging over even longer and more varied foraging paths consistent with observations of foraging trajectories of mice in the wild [51].

Whilst AQUA makes decisions to stay or leave, it does not use time elapsed in patch or a calculation of reward rate (necessitating the estimation of timing information) in order to make its decisions. This is a fundamental aspect of MVT and other OFTs, and indeed recent work suggests that midbrain DA has access to this alongside a second axis of value in such a way that could be beneficial for patch foraging decisions [52]. Here we provide evidence a) that untrained mice do not time their visits *per se* (Supp. Fig. 5) - yet still achieve remarkably high reward rates and b) that reward rate estimation is not necessary to produce the same predictions as MVT even in an environment where patches deplete exponentially (Supp. Fig. 13). However, it will be interesting in future work to explore whether and when time estimation emerges as a necessary component to foraging with more experience, particularly considering its importance for some theories in associative learning [53]

The success of AQUA in accounting for both moment-to-moment choices and variability across mice inspired us to use it as a tool for interpreting observed dopamine signals. There are two algebraically equivalent computations that can implement the core update rule of AQUA. This presented us with an opportunity to compare which, if any, of these computations were consistent with observed dopamine responses to reward delivery. Our simultaneous photometry recordings of dopamine responses to reward in the core of the nucleus accumbens (NAcC) were uniquely consistent with the global dynamic learning rate component of AQUA. Whilst some aspects of NAcC responses are consistent with RPE - in particular, the difference in response between rewarded and unrewarded visits (though that is not a strict test of RPE *per se*) - there were also critical inconsistencies with RPE especially pronounced around changes in option quality. In the context of FoMO a dynamic learning rate should start high around session beginning and slowly relax over time. We are not the first to observe such a decay of reward responses within a session in a manner consistent with a dynamic learning rate; recent re-analysis of existing datasets have detected such trends that were missed previously [54]. In addition, here we also show that re-analysis of dopamine signals in NAcC from a recent study in well-trained rats moving between three reward options also exhibits similar reduction of reward responses across at least as many datasets where RPE correlates are apparent [22] (Supp. Fig. 12). It is clear that animals rapidly and robustly learn the quality across all 6 reward options and rapidly adapt their behavioral policies across multiple changes in port quality arrangements over the first 5 sessions in the absence of obvious RPE coding in NAcC dopamine signals. Therefore, this work also emphasizes that robust reinforcement learning can occur in the absence of detectable RPE correlates in midbrain dopamine neurons consistent with observations of early learning in other paradigms as well. Future developments of the modeling approach taken here could integrate further complexity in the dynamics of the global (shared by all options) learning rate term and or include additional local (option specific prediction error) modulation as perhaps emerging with more extended training. Indeed, our re-analysis suggests both components may be present in NAcC dopamine signals with more extended training (Supp. Fig. 12).

Global parameters that govern how RL agents execute policies are, in fact, a component of many models. For example, the balance between explore/exploit can be thought of as a global signal. A global modulation of learning rate provides an attractive addition that can accomplish at least some of the desirable properties of explore/exploit state switches such as allowing rapid adaptation of agent policies when environmental contingencies change. Future work will be critical to understand how to best combine additional global modulation signals, perhaps drawing inspiration from models in machine learning that exploit multiple such components to efficiently train models. Dopamine signals in the dorsomedial striatum (DMS), in contrast to those in NAcC, were neither consistent with RPE nor clearly consistent with a global learning rate, highlighting an emerging trend towards observing subtype dependent responses in the heterogeneous midbrain dopamine neuron population. DMS dopamine may be more critical for modulating aspects of the movements that underlie the behavioral demands of traveling between ports, but that remains to be examined in detail.

## Methods

### Animals

All protocols and animal handling procedures were carried out in strict accords with protocol 22- 0234.03 approved by the Janelia Institutional Animal Care and Use Committee and in compliance with the standards set forth by the Association for Assessment and Accreditation of Laboratory Animal Care (IACUC). Data from a total of 23 mice are included here. This includes two main cohorts: 17 *Slc6a3* (DAT-cre) for use with GCaMP8m and 6 C57BL6 mice for use with dLight (see *Photometry*). All mice were aged between 3 months and 1 year at the time of experiments. 20 mice experienced configuration 1 (Fig. 1c), and all photometry data is from this subset of mice.

Mice were singly housed on a free-standing, individually ventilated (about 60 air changes hourly) rack (Allentown Inc.). The holding room was ventilated with 100% outside filtered air with >15 air changes hourly. Each ventilated cage (Allentown) was provided with corncob bedding (Shepard Specialty Papers), at least 8g of nesting material (Bed-r’Nest, The Andersons) and a red mouse tunnel (Bio-Serv). Mice were maintained on a 12:12-h (8am-8pm) light/dark cycle and recordings were made between 12pm and 7pm. The holding room temperature was maintained at 21+/-1C with a relative humidity of 30% to 70%. Irradiated rodent laboratory chow (LabDiet 5053) was provided *ad libitum*. Following at least four days’ recovery from headcap implantation surgery, animals’ water consumption was restricted to 2ml per day for at least three days before training. Mice underwent daily health checks, and water restriction was eased if mice fell below 75% of their original body weight.

### Arena hardware and software

#### Main arena construction

The main body of the arena (Supp. Fig. 1) was constructed from stainless steel with interior dimensions (LxWxH) 83.75in x 17.38in x 22.50in. Walls were formed from 0.063” thick 304 stainless steel. For ease of autoclaving, the arena was composed of three separable subassemblies fastened together with custom-manufactured latch assemblies consisting of a lift-off hinge, for alignment between subassemblies, and draw latch for a secure hold. Panel inserts were used to reduce reflections from the walls. These were fabricated from matte white acrylic sheets and laser cut on an EpilogLaser Fusion Pro 48 laser engraver. These inserts were held against the walls of the arena using a pair of loop clamps hooked over the arena top edge, while end panels are held in place by the presence of side panels. Permanent magnets at the spout locations were also installed using 3D printed ‘plugs’ to ensure the panels remained flush against the arena walls. The plugs were printed on a Stratasys Connex 350 PolyJet printer from VeroWhitePlus resin. The whole assembly, including main body of the arena and subsequently described frame, was designed in Autodesk Inventor and parts printed through Stratasys’s GrabCAD Print and Objet Studio slicers.

To raise the arena above ground and for installation of the tracking camera, a frame was assembled predominantly from 1.5” aluminum T-slot extrusions of exterior dimensions (LxWxH): 0.99in x 27.00in x 90.00in. The frame consisted of two subassemblies - arena-supporting and moving-stage supporting - in order to independently support the body of the arena. This separation eliminated the transfer of vibration from the moving stage to the arena. The stage-supporting subassembly featured a 2250m-long LC40 Zaber stage, on which was mounted an AB104 Zaber 90 degree angle bracket for ease of mounting recording equipment.

#### Moving gantry tracking camera system

We built a bespoke tracking camera system in order to follow animals’ trajectories as they moved around the arena. This also allowed for movement of a commutator for photometry recordings or other hardware to be kept close to the animal, permitting mice to move unperturbed across the full area of the arena whilst recordings took place. The main components of this system were a high speed camera mounted onto a stand attached to a stage that moved in close-loop with the mouse. We describe both the hardware and software implementation in detail below.

##### Hardware

We used a high frame rate camera (Basler ace acA2040-180km NIR) and camera lens (Edmund optics 12mm-36mm focal length, varifocal video lens) to record mice as they moved around in the arena. Video recording was done in an environment where the illumination wavelength is out of the mouse visible spectrum (>700 nm). To achieve this, we mounted an 850nm IR LED ring light (NorthCoast Technical) with intensity control to the camera. The camera set up (as well as a custom-designed 3D printed commutator holder) was mounted onto a motorized moving stage (Zaber Technologies) that itself was mounted onto the aforementioned frame that surrounded the arena. The stage had a maximum of 2250mm travel, with integrated controller and motor encoder for recording of stage position, and was able to achieve a maximum speed of 3m/s. The camera’s field-of-view was chosen to be greater than 18 x 18 inches, so that any movement of animals along the Y-axis does not require the camera to be moved. That is, the entire width of the arena is visible at any given time such that the animal needed to be followed only along the X-axis.

Images were captured every 5ms and processed in real time using an FPGA module NI PCIe-1477. Images from the camera were transferred directly to the FPGA via dual high speed camera link cables. Camera link cables are capable of transferring data up to 850 MB/s. The FPGA module PCIe-1477 incorporates Xilinx Kintex-7 325T FPGA and was installed in a PCIe slot of a Dell precision 5280 workstation. FPGA processed images were then transferred to a custom-made LabVIEW software module and finally data were saved in a solid-state hard drive for data analysis. Zaber stages were controlled via USB-UART interface adapter for sending motion commands and receiving stage position read out.

##### Software

We used the software tool LabVIEW to achieve high speed video recording and real-time animal trajectory tracking. A GUI was also built in LabView for easy user adjustment of recordings. With our custom developed FPGA logic, we have achieved animal position identification in real time via image processing with a latency of sub-millisecond. Identified animal position was fed back to the Zaber stage to follow animal trajectory. The image processing algorithm consisted of three steps:

1. Converting grayscale image to binary image at some user defined threshold range.
2. Separation of the binary image into multiple objects based on connectivity.
3. Application of a filter parameter, area. Here Area defines the number of pixels present in the object, in this case the mouse.

Videos were captured at 1000 x 1000 pixels at 200 fps and saved in binary format (.bin) directly to a solid-state drive to achieve the highest possible writing speed, and were subsequently converted from .bin to .avi for image analysis. Information recorded by the system included the current location of the mouse within the field of view of the camera in X and Y coordinates. The saved video was cropped around the identified largest object (the mouse) to reduce file size (640 x 640 pixels). The crop location was maintained at a fixed distance from the recorded X/Y location of the mouse, and so obtain the absolute X/Y position we performed the following transforms after interpolation between points:

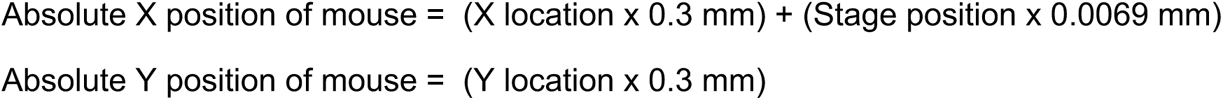

For subsequent visualization and analyses, these values were then smoothed across a mean rolling window of 20 samples.

#### Lick spouts

Six custom made lick spouts were mounted onto the walls of the arena using a magnetic ‘plug’ system. Each spout consisted of a metal lick tube (ID 1.5mm, OD 2mm) that extruded into the arena and that mice had access to, connected to tubing (Cole-Parmer) linked to a 24V solenoid (Lee Company) that was activated upon capacitive touch (Sparkfun Capacitive Breakout) if reward was available. An independent 5ml reservoir provided water per spout and spouts were calibrated regularly to ensure a consistent amount of water delivery. All elements of the spout - reservoir, capacitive sensor, solenoid, and metal lick tube - were mounted on a custom designed 3D printed holder, designed in OnShape and printed on a Stratasys F170 FDM printer from ABS filament. The capacitive sensor also interfaced with a pyControl board [55] used to receive sync pulses and record behavioral input as subsequently described.

#### Additional arena miscellany

During experiments the arena floor was covered with 1-2 inches of Alpha Dri bedding (Lab Supply). This had several benefits: mice were more comfortable running at speed across this surface in comparison to a smooth floor, and the white color of the bedding provided clear contrast to the black mouse for more accurate tracking by the moving camera. The bedding was refreshed between every mouse to ensure less carry over of scent across animals. Aside from bedding we also included a laser cut red acrylic shelter in two parts that was secured using magnets onto the floor in the center of the arena, and a pellet of standard lab chow was also placed nearby.

All experiments were conducted in semi-darkness. On day one, mice were completely naive and had no pre-exposure to the arena environment. Because of this the metal tubes of the spouts were each baited with a small drop of water to allow mice to understand that they could retrieve water from these locations through licking. Aside from this initial baiting, no guidance or cues were given to mice to indicate reward availability. pyControl was used for task programming and interfacing with both the lick spouts and the tracking camera. Mice were assigned to one of two configurations (Fig. 1c) and a subset of mice on configuration 1 experienced a 180 degree rotated version.

### Fiber photometry

#### Surgery

Slc6a3 mice (n = 14 in final dataset) were intracranially injected with AAV2/1-syn-FLEX-jGCaMP8m-WPRE, 2/1-CAG-flex-rev-tdtomato (Janelia viral core) and sterile saline mixed in 2:1:2 ratio at a volume of 400nL. Mice received injections bilaterally, in the VTA (-3.3A/P, ±0.4M/L, -4.3D/V) and in the SNc (-3.0A/P, ±1.4M/L, -1.4D/V). Custom 0.39NA 200um diameter fiber cannulas were implanted respectively in the NAc (+1.4A/P, ±0.8M/L, -4.1D/V) and DMS (+1A/P, ±1.7M/L, -2.3D/V) and the fiber with the best signal was chosen for subsequent recordings.

C57BL6 mice (n = 6 in final dataset) were injected with undiluted AAV5-CAG-dLight1.3 and AAV5-hsyn-tdTomato at a volume of 200nl. These mice also received injections bilaterally, in the NAc (+1.4A/P, ±0.9M/L, -4.1D/V) and DMS (+1A/P, ±1.8M/L, -2.4D/V) with fiber cannula implantation at the same sites. Again, the best signal was chosen for recordings.

#### Photometry recordings

Dopamine calcium activity (GCaMP) and release (dLight) were recorded at a sampling rate of 130 hz using pyPhotometry [56]. The optical system comprised 465 nm and 560 nm connectorized LEDs, a five port minicube with integrated photoreceivers, and a pig-tailed fiber-optic rotary joint (Doric Lenses). Time division illumination with background subtraction was applied to prevent cross-talk between fluorophores due to overlapping emission spectra and/or ambient light. The photometry signal was aligned to behavior recorded by the pyControl system through sync pulses received by a digital input of the photometry board.

#### Histology

Mice were terminally anesthetized (isoflurane, >3%) and perfused with ice-cold phosphate buffered saline, followed by paraformaldehyde (4%wt/vol in PBS). Brains were post-fixed for 2h at 4 degrees C and then rinsed in saline. Whole brains were then sectioned to 50 um thickness using a vibrating microtome (VT-1200, Leica Microsystems). Sections were imaged using an Olympus MVX10 dissection microscope and fiber tip positions were estimated by referencing standard mouse brain coordinates [57]; Supp. Fig. 8).

### Analysis

All analysis was performed using custom Python and MATLAB code.

#### Behavior analysis

##### Preprocessing of choice data

Lick data at each option consisted of a train of ‘lick on’ and ‘lick off’ events as the animal’s tongue touched and then moved away from the metal spout tube. We used this to clean the data by first ensuring that each ‘on’ event is followed by an ‘off’ event. Additional visualization including inter-visit interval distribution was used to ensure that spouts were not erroneously activated. Once confident that the licks captured were accurate, visits were then identified as either requiring a switch from one option to another - which was not thresholded by inter-visit interval - or >1s between licks at the same option.

##### Multinomial logistic regression

We developed a multinomial logistic regression approach in order to predict decisions across six options based on a number of different predictors. Note that use of a multinomial approach allowed for estimation of the probability of a given choice across a range of six options, i.e. an extension of binary logistic regression. A predictor matrix across choices for each mouse and session was generated, of size *n* features (predictors) by X *c* choices. This was then used in training a regularized estimated coefficient tensor of size *k* X *n* X *c* on 80% of the data, where k is the number of options (six). Through softmax transform this resulted in a *k* X *n* probability matrix, and argmax (greatest value) was used to identify a single predicted decision and compared to the true labels iteratively to jointly optimize the tensor through loss function minimization. The beta values from this learned coefficient tensor is what is plotted in Fig. 4g (left). These values were also used on the remaining 20% (test) dataset for 5-fold cross-validated estimate of accuracy in Fig. 4g (right). All predictors were normalized between -1 and 1.

#### Photometry analysis

##### Preprocessing

Photometry signals were preprocessed using custom Python code. In brief this involved the following steps:

1. Lowpass filtering
2. Baseline fit of the signal (F) using a double exponential where possible
3. Highpass filtering
4. Movement correction of the signal through linear fit + subtraction from control channel (delta F)
5. Bleaching correction by dividing F by delta F

A full outline and example of these steps is contained in the following jupyter notebook: <u>fiber</u> <u>photometry preprocessing</u>

##### Photometry regression

To visualize the change in coefficient weightings across time as in Fig. 5d, a ridge linear regression was run on each timepoint to predict dopamine activity. For the full regression across both rewarded and unrewarded visits, predictors included whether the visit was rewarded (0,1), option quality (6:1), whether the previous visit was rewarded (0,1), whether the previous choice was at the same option (0,1), the interaction between previous choice and reward (0,1), the quality of the previous option (6:1), whether the visit occurred before or after the random period (0,1), time since the last visit at the option, and distance between current option and previous option. The reward-only regression in Fig. 5f included the additional predictors of AQUA alpha value and AQUA RPE value, which were both updated on a visit by visit basis. All predictors were normalized between -1 and 1 and mean subtracted before being z-scored allow direct comparison of predictor weightings.

### Modeling

#### Static model

As a benchmark for animal performance, we used generative modeling to simulate three static choice policies: random, omniscient, and heuristic win-stay lose-shift (WSLS). We termed these policies ‘static’ as they do not change with learning. On a session by session basis, the model received the timing of animals’ sampling across all options but without information as to which option was chosen at each sample. The model then chose what option to sample from based on the given policy. Whether the choice is rewarded depends on the empirical timings of the reward intervals in the task, matching animals’ experience.

The choice policies work as follows: in the case of the random policy, all options were chosen with an equal probability. In WSLS, the model kept track of the outcome of the previous choice. If the previous choice was rewarded, the model repeated the same choice again at the next choice timepoint. If the previous choice was unrewarded, the model randomly selected from all other remaining options at the next choice timepoint. The ideal agent knew when reward is available at each of the options and selects an option that has reward available whenever possible. If multiple options are baited at that timestep, the agent chooses the one associated with the worst option. If no ports are baited, the agent randomly selects across all ports. Note: despite this level of ‘omniscient’ knowledge, the omniscient agent will not retrieve all available rewards in the task across port schedules as it is limited by the animals’ empirical sampling rate. The omniscient agent therefore provides an upper limit to the number of rewards an animal in a given session could in theory have collected considering the frequency with which it sampled the ports.

We used several approaches to compare mouse behavior to different simulated static policies. To quantify the moment of deviation from random policy in cumulative reward, we calculated the difference between the slope of lines fitted to moving windows of 10 visits for both animal and averaged random agent behavior from 100 simulated runs. A slope difference threshold of 0.02 was chosen as this consistently accurately identified divergence. To understand which simulated policy was most similar to animals’ behavior, we calculated the ratio of KL divergence.

#### Optimality model

To obtain the optimality model here, we assume that the policy to be optimized is the 6 x 6 set of possible transitions in FoMO. We constructed a generative simulation where decisions are made based on this transition matrix. After each choice, the time to travel to the next port is drawn from the animal’s empirical distribution of travel times between those specific options. Upon arrival at a port, the duration of the visit is also sampled from the animal’s empirical distribution of visit durations.

This simulation allows us to calculate the average reward rate (number of rewards/time) for any given set of transition probabilities. Because of the stochasticity of the decisions, we also average reward rates over 10 simulation trials to get the overall mean reward rate. To find the optimal transition matrix, we begin with an initial guess for the parameters - either a uniform matrix (equal probabilities of visiting each port) or estimated probabilities from behavioral data. We then iteratively update the parameters using MATLAB’s ‘patternsearch’ function until the solution converges, meaning that further perturbations do not improve the mean reward rate. This is treated as the optimal solution, and the optimal model corresponds to the generative simulation using this policy.

#### AQUA Learning model

In the development of a computational model to simulate reinforcement learning agents that allocate behavior across a set of 6 options (generalizes to an arbitrary number), we considered a number of different update algorithms that could be used. The full model newly developed for this study we named AQUA: **A**daptive rate **Q**-learning of **U**tilities for **A**pportioning actions. As the name suggests we combine 4 critical features that are based upon insights from machine learning, reinforcement learning and foraging theory. The critical features are:

1. Maintaining a separate estimate for the probability of reward given the agent stays at the current option. This is analogous to the stay/leave decision central to foraging theory and a core feature of MVT. The update equation is the same as described in eq. 1 below, but specific to transitions from ***O_N_ > O_N_***.

2. Updating a running estimate of probability of reward for each of the 6 options if transitioning from any other option. This leads to a 6 place vector ***P(R|O_N_)*** for each option ***N***. Estimate updates can be thought of as a calculation of a running sample mean of a binary variable ***R={0,1}.*** The estimated probability is thus increased by an increment proportional the learning rate ***α*(*τ*)** on rewarded visits balanced against a memory term proportional to **1-*α*(*τ*).** For this reason ***P(R|O_N_)_t_*** is akin to an estimate with a window size ∼***1/α*(*τ*):**

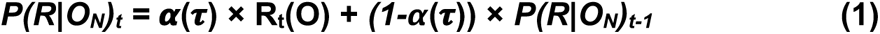

We note that this equation can be recast in an algebraically equivalent form that introduces a prediction error term, namely:

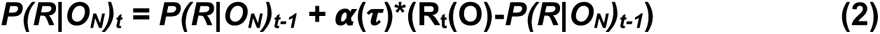

Thus, the update equation bears strong similarities to the update equation from Q learning (hence the Q in AQUA).

Note that in typical formulations reward (R_t_) would be defined as {0,1}. However, this needn’t be the case and other possibilities could be considered depending upon implementation in biological circuits. For example, if one defined an unrewarded visit as, say, -0.1 rather than 0 this will introduce a bias into the estimate provided by P(R|O_N_) relative to the true probability. The extent of the bias can be mitigated by reducing the learning rate term on unrewarded trials down to a smaller value, say 10%. In this case, due to the noisiness of small sample sizes in a 3 hour session as in FoMO one can readily confirm with simulation that an essentially undetectable change in the estimate of P(R|O_N_) is predicted. Further note that a bias is not necessarily a bad thing in certain situations. For example, a downward bias in a reward probability estimate can allow a more rapid abandonment of an option with a rate determined by ***α*(*τ*).**

This is one interpretation of how decrements in the NAcC dopamine release specifically on unrewarded trials can be consistent with AQUA. We do not consider this formulation in detail because it adds additional parameters and complication, but do note that there is nothing inconsistent in principle between the observed unrewarded NAcC responses and a learning rate interpretation of NAcC dopamine activity. Moreover, as noted in the main text, recent work indicates that suppression of dopamine responses on unrewarded trials is most consistent with mediating a decreased learning rate [43].

3. Where the learning rate ***α*(*τ*)** is a decaying function of total visits (***τ***) shared by all options - i.e. a “global” learning rate. ***α*(*τ*)** decays exponentially with a time constant ***α*_*τ*_** with a peak value of ***α*_peak_** and an offset ***α*_offset_**. All of these values are free parameters used in fitting the model to behavior. Optimization was evaluated by maximizing an objective that jointly maximized prediction of mean choice behavior (transition matrix correlation) and minimized root mean squared error in income (smoothed rewards per visit as a function of time): Obj = rho_transitions_ − rmse_income_. This could be differentially weighted on each term, but simulation data reflects an event weighting. A relaxing learning rate here, given a prior of equal P(R|O) for all options, is equivalent to relaxing from random exploration - equivalent to a feature often exploited in machine learning. For example, it was used to great effect in the (effectively Q learning) agents trained to play Atari games [37] and in such cases has been interpreted as similar to a global property often referred to as “explore/exploit”. More generally, dynamic learning rates that effectively relax from a large to small final value are now ubiquitous in the use of ADAM in machine learning and artificial intelligence [36].

4. Rather than simply computing the value of each option (probability of reward) we also use a distance weighting term where the pairwise distance between the current option and a distal option is governed by the geometry of the environment according to a Fitts’ Law like function distance cost:

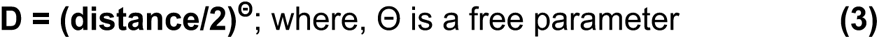

We estimate this free parameter by scanning over a range 0.5-3 fit to the most stable period of behavior (end of session 2). Most animals were best fit by a Θ∼1.5. The distance cost is used to scale option quality:

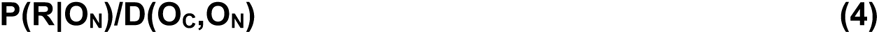

which is then passed to a proportional sampler to render a choice. Because the value of an option is scaled by a cost term we describe these as utilities. The analogy to Fitts’ Law reflects an implicit conjecture in the model that distance weighting may not be an effort cost as often presumed, but perhaps more akin to a navigational cost associated with a more general notion of the increasing challenge of accurately navigating to a distant target. This will be the subject of future work, but at this point is an empirical observation that nonlinear distance costs prohibited modest, but reliable, improvements in explanatory power.

Given the above overview of AQUA, we next describe the structure of the simulations and briefly touch upon the algorithms used to simulate a number of standard agent models for comparison.

Environment Setup: The model initializes an environment with the reward schedule based on the empirical data from a given animal’s sessions. It sets up the port intervals and reward probabilities for each port at each time step. For a given simulation at each timestep we use the data from a given animal to decide whether a visit is attempted. The effective sampling rate of the simulations is 1Hz.

Simulation Loop: For each time step, the model follows these steps:

a. Determine if the subject should make a choice based on the sampling schedule.
b. If a choice is to be made, decide whether to stay at the last checked port or switch based on the stay probability.
c. If switching, select a port based on the estimated option quality in proportion to 1-ε. In the case of AQUA this decision is the option quality normalized by the distance cost. In the vanilla Q learning baseline the option quality is the value estimate passed through a softmax function with beta=1. Random agents or WSLS agents make a random choice. Finally, the “ideal” agent chooses randomly amongst the options that have an available reward (or randomly if no options have a reward available). For all agents a noisy random choice was made with p=ε. A range of values for ε were tested, but in general most simulations used ε=0.1.
d. Check if the chosen port provides a reward (R(O_N_)=1) based on the ground truth reward schedule.
e. Update the agent’s P(R|O_N_) according to the update equation specific to the agent model.
f. Record the subject’s choice, reward, and other relevant variables.

Evaluation: The model computes various performance metrics, including:

- Cumulative rewards.
- Local income: the average reward rate within a sliding window.
- Total visits to each port.
- Transition probabilities between all pairs of options for each sequential visit pair.

---

**Supp. Fig. 1.**
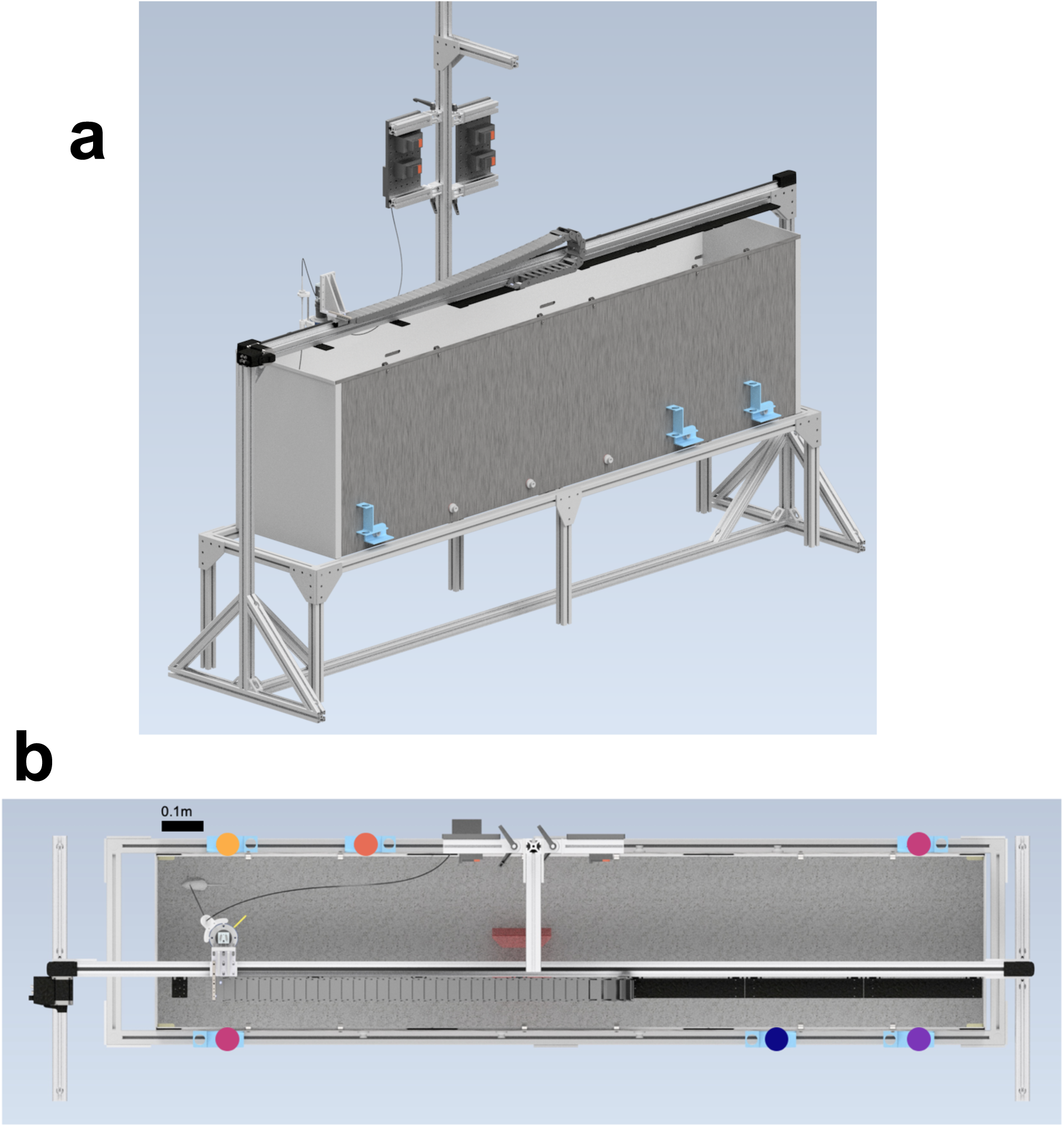
**A** CAD model of FoMO arena. **B** Top-down view of CAD model, with example arrangement of option quality.

**Supp. Fig. 2.**
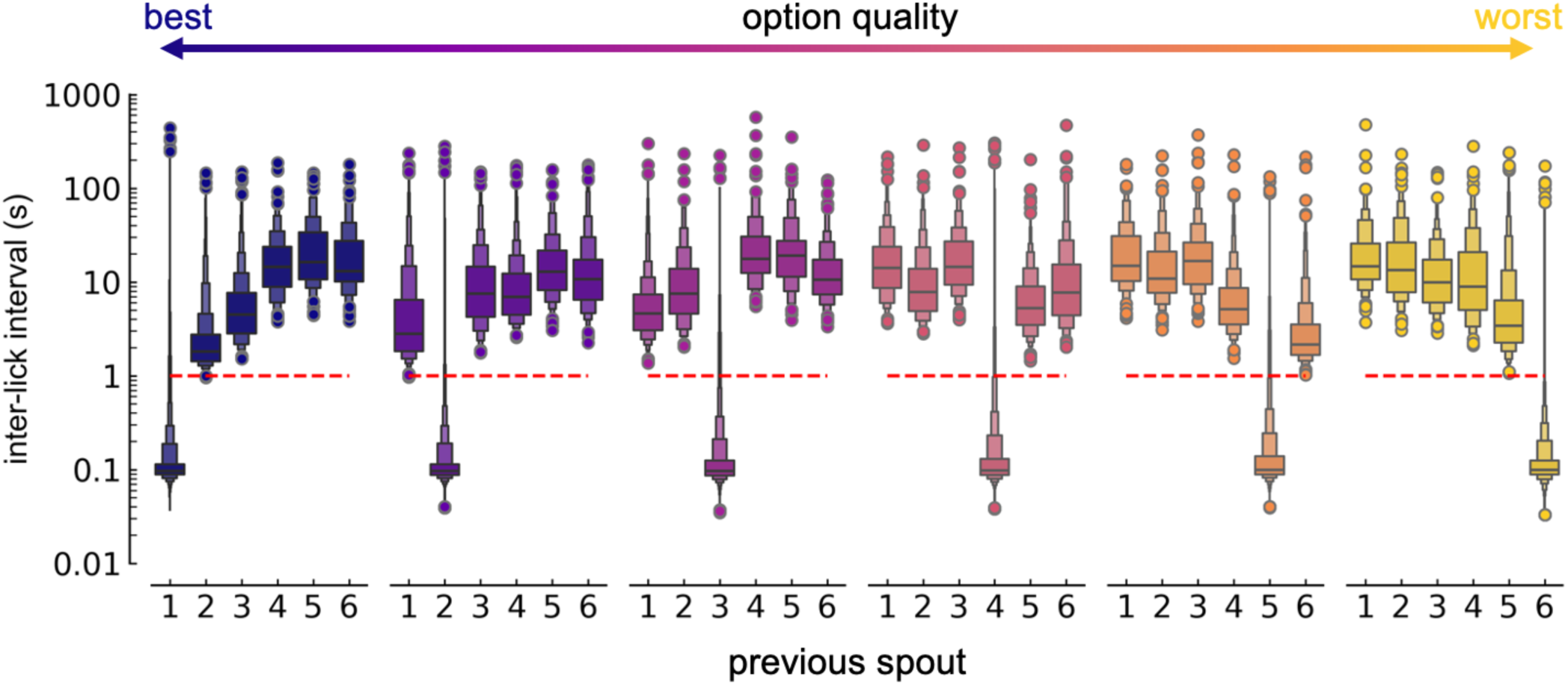
Identifying a threshold for defining option ‘visits’. Raw unaveraged inter-lick interval distributions from all animals shown. Red dotted line indicates threshold which was set at 1s: licks occurring at a latency greater than 1s were classified as separate visits even if to the same spout.

**Supp. Fig. 3.**
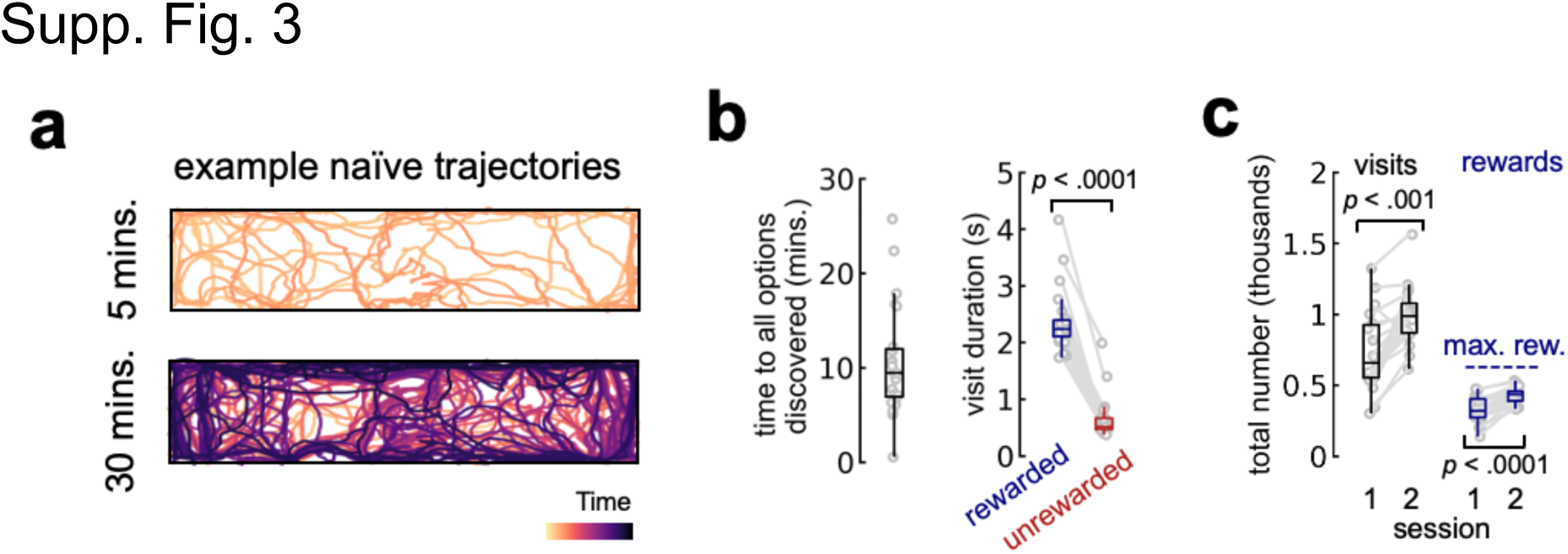
**A** Trajectories from an example mouse within the first five minutes (top) and 30 minutes (bottom) of the first session. **B** Left: time to all options discovered. Right: mean visit duration for rewarded and unrewarded visits. **C** Total number of visits (left) and rewards) right for sessions 1 and 2. Box plots represent median at center bounded by 25th and 75th percentiles of the data, with whiskers extending to the extent of the distribution barring outliers. All dots represent individual mice. Significance testing: non-parametric Wilcoxon rank sum (f, g, k).

**Supp. Fig. 4.**
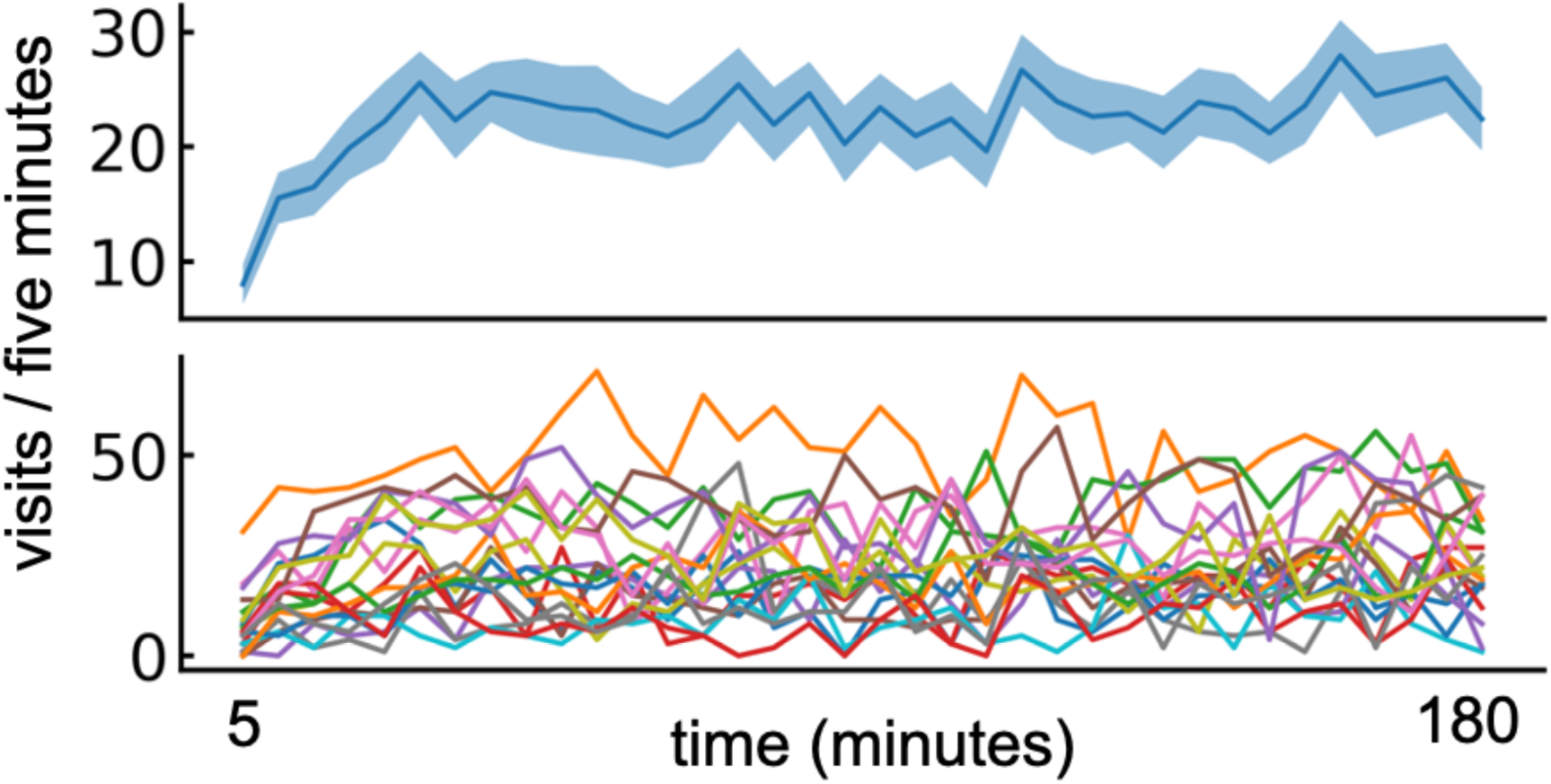
Visit rate across all ports remains stable across the first session indicating that mice are engaged throughout. Top: mean and SEM of number of visits per 5 minute, non-overlapping bins. Bottom: same as top but one line per mouse.

**Supp. Fig. 5.**
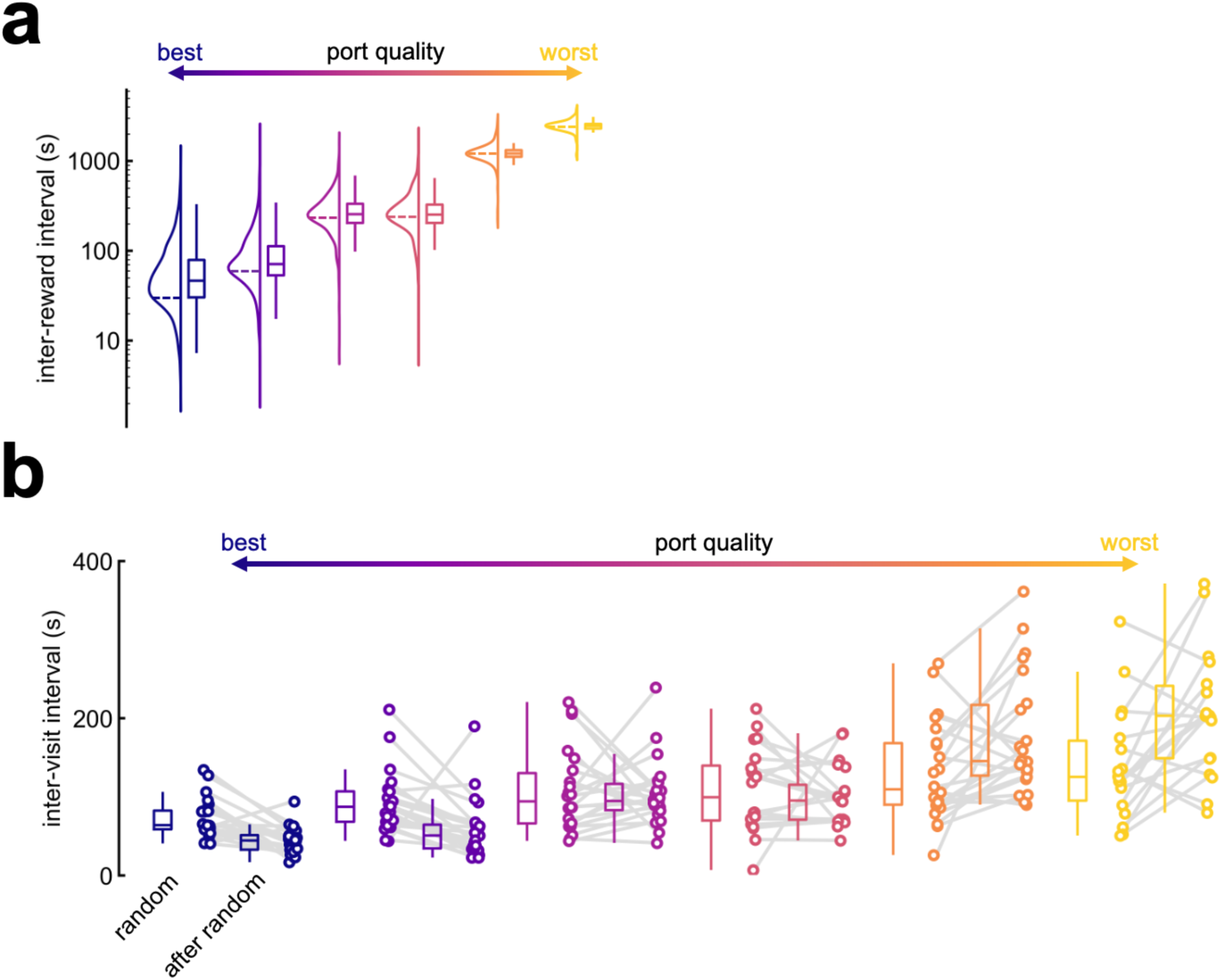
**A** Inter-reward interval by option concatenated across all animals. For each option, on the right the violin plot shows the full distribution and on the left the box plot shows quartiles with center as median. The dotted line crossing the violin plot is positioned at the schedule’s true interval (i.e. 30s, 60s, etc.) **B** Inter-visit interval by option split by before or after an individual animal has deviated from random (calculated as shown previously). For each option, on the left the box plot shows quartiles with center as median, and on the right individual points represent the mean for each animal.

**Supp. Fig. 6.**
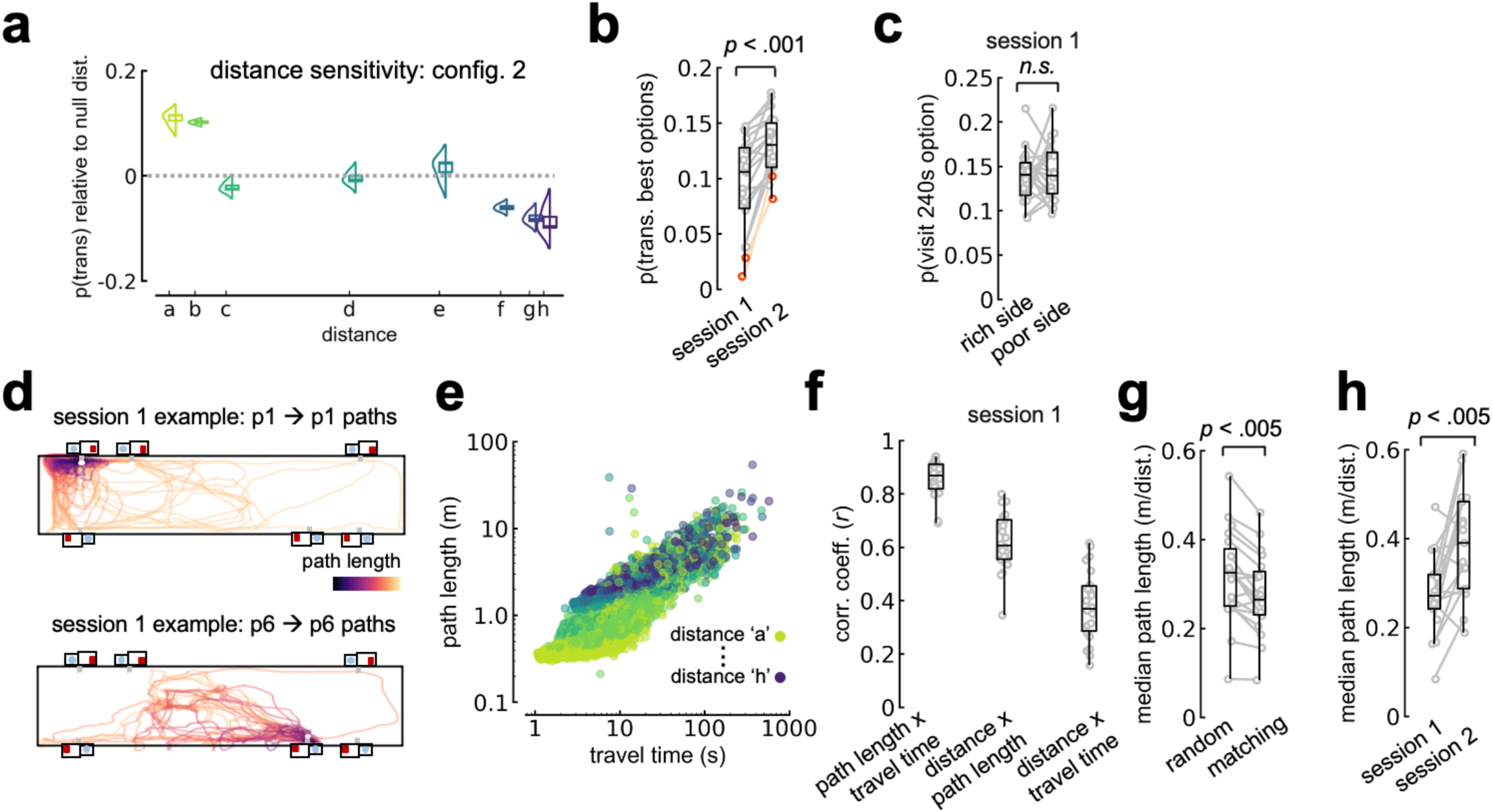
**A** Probability of making a transition by pairwise distance, relative to null distribution (overall probability of making the transition against all transitions) for configuration 2 mice only. **B** Probability of transitioning between the best and second best options by session. **C** Probability of visiting the 240s option by whether it is grouped with the good (‘rich’: 30 and 60s options) or bad (‘poor’: 1200 and 2400s options) quality alternatives. **D** Example return paths to the best (top) or worst (bottom) options. **E** Left: path length by travel time. Dots colored by pairwise distance. **F** Spearman’s correlation between path length and travel time (left), distance and path length (middle), and distance and travel time (right). **G** Median path length travelled by mice during random or matching (after random) behaviour in session 1, normalized by objective distance. **H** Median path length travelled by mice in session 1 and session 2, normalized by objective distance. In all plots each dot is one mouse, aside from F where each dot is a single trajectory. Box plots represent median at center bounded by 25th and 75th percentiles of the data, with whiskers extending to the rest of the distribution aside from outliers. All significance testing: non-parametric Wilcoxon rank sum. ***See Appendix 1 for discussion of this figure.***

**Supp. Fig. 7.**
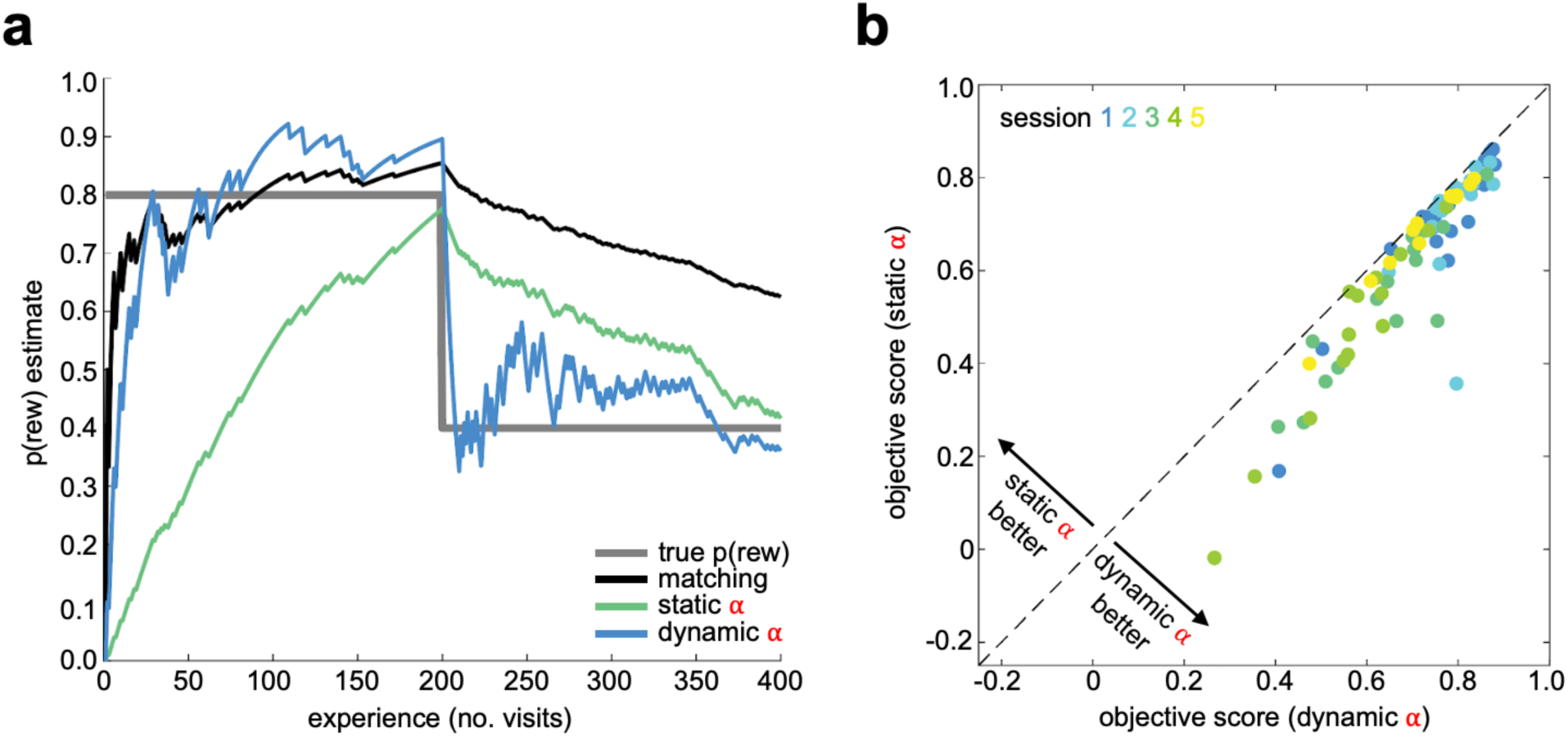
**A** P(rew) estimate for one example sequence of rewarded/unrewarded visits. True generative probability is 0.8 for first 200 trials and 0.4 for last 200 trials. We consider several different update rules for an estimator: a straight matching estimator that divides the full history of collected rewards by the number of visits (black), the P(R|O) update equation from AQUA using either a static alpha value (0.01 but similar results hold for a range of values; green) or a dynamic alpha that relaxes to the same static value (blue). **B** Objective score (explained variance normalized by RMSE income as defined in Methods) to compare fit quality between AQUA simulation and observed behaviour for static vs. dynamic alpha across all 5 sessions (including schedule changes as described in fig. 6). In all cases dynamic alpha provides a better fit (p<<<0.01).

**Supp. Fig. 8.**
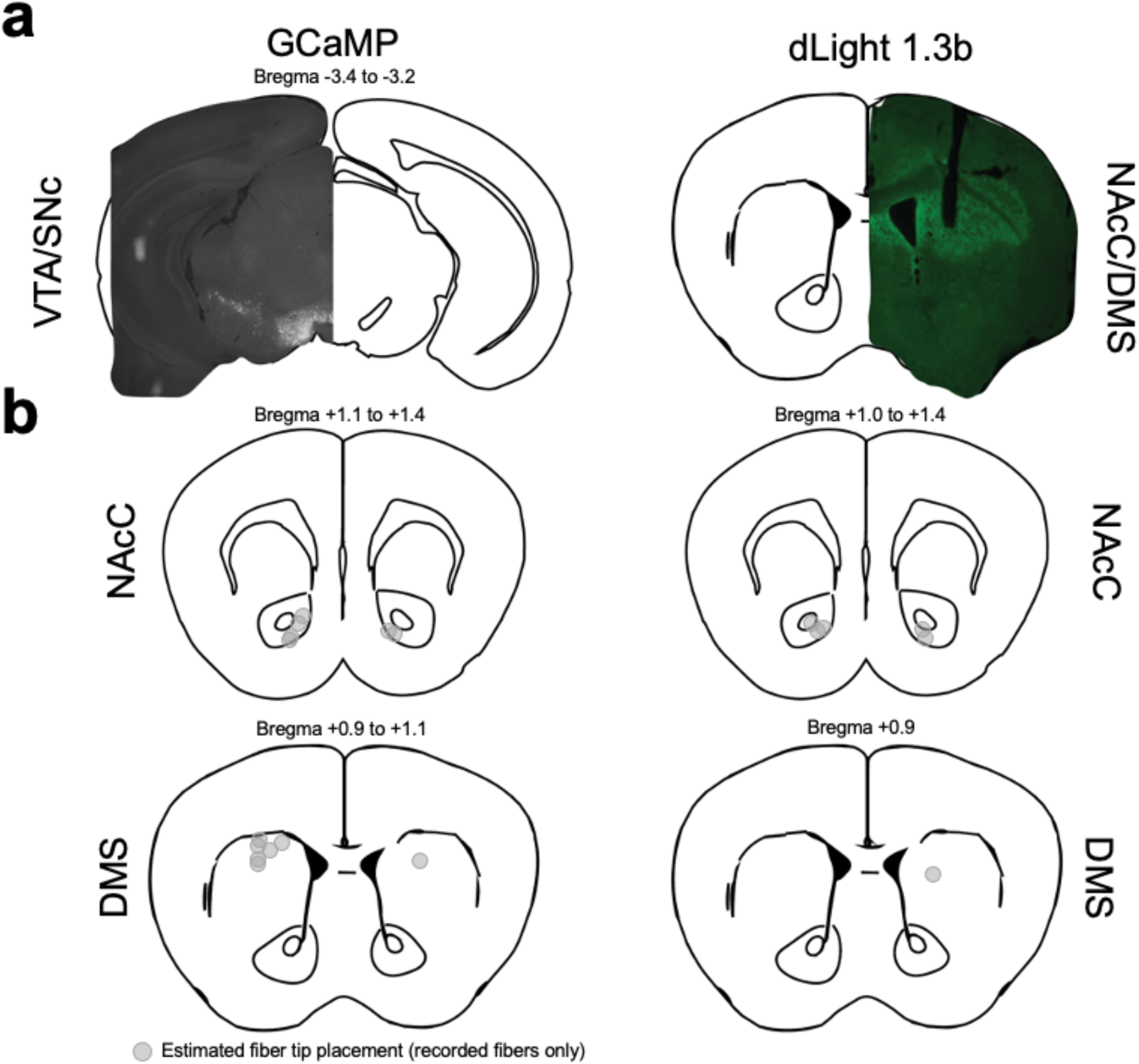
**A** Example histological verification for GCaMP injections in VTA and SNc (left, VTA visualized) or dLight 1.3 injections in NAcC or DMS (right, DMS visualized). **B** Estimated fiber tip placement for GCaMP (left) and dLight (right) recordings. Adapted from Paxinos & Watson, 2019.

**Supp. Fig. 9.**
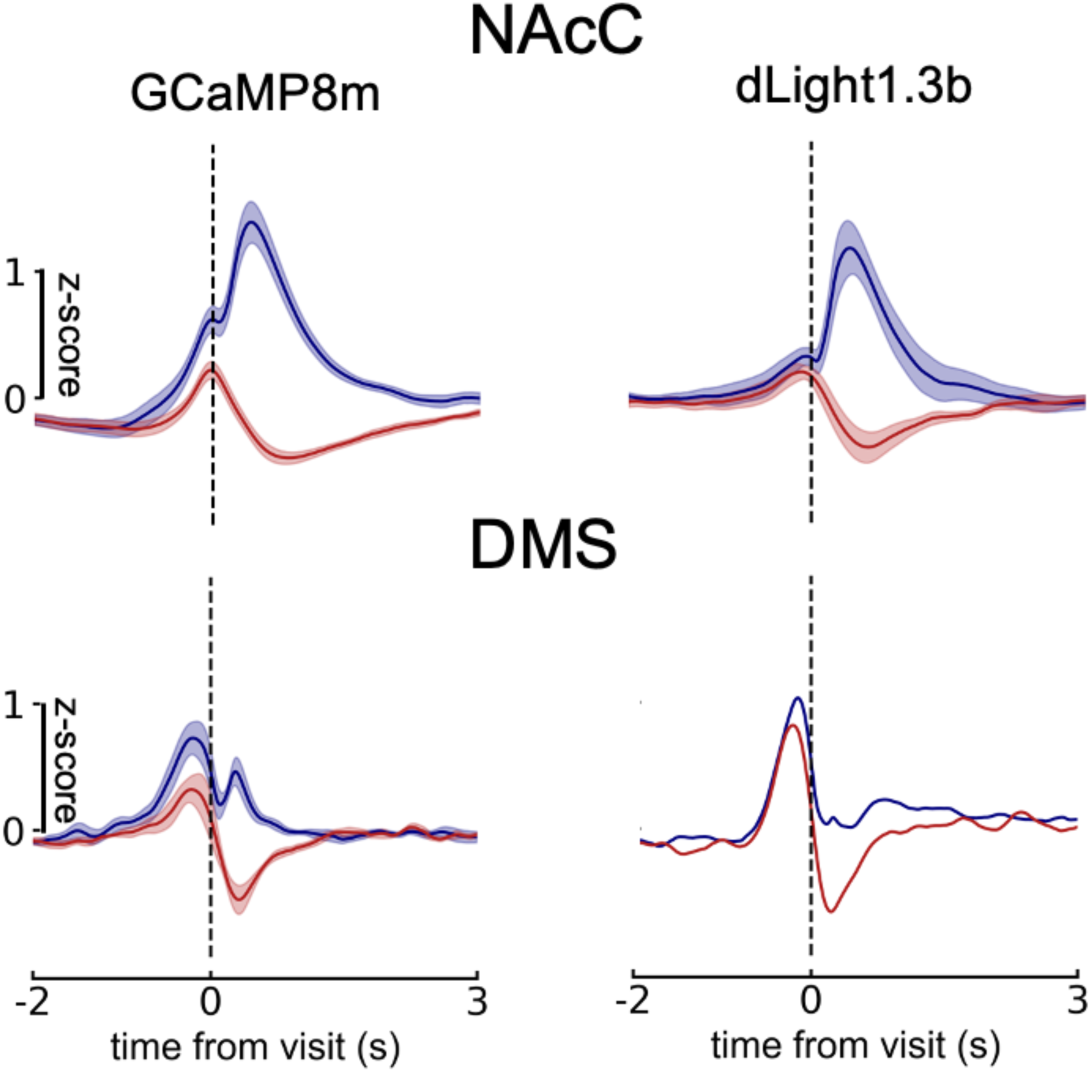
Responses to rewarded (blue) and unrewarded (red) visits, split by sensor and region. Solid line is mean, shaded area is SEM.

**Supp. Fig. 10.**
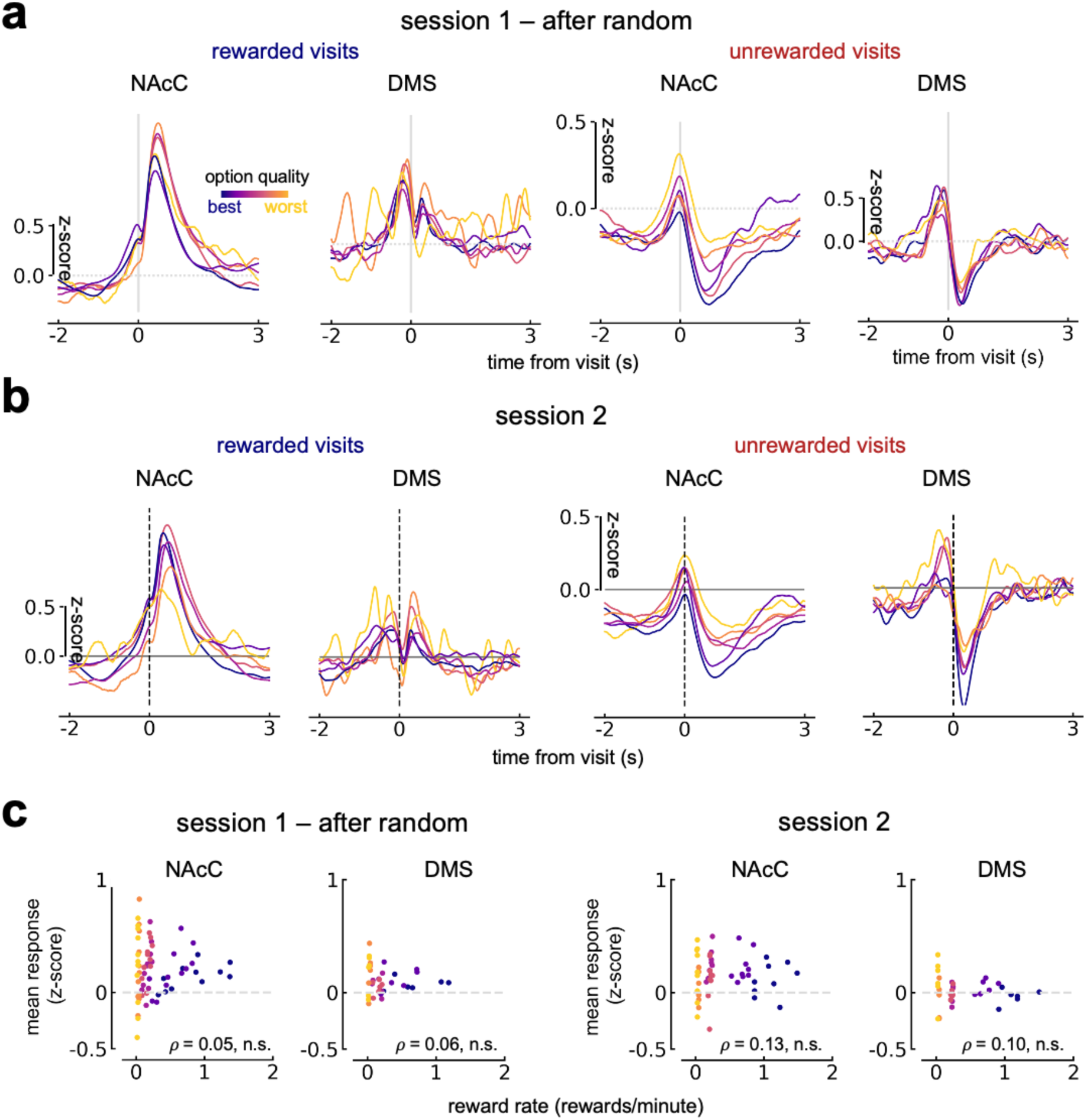
**A** Mean responses to rewarded (left) or unrewarded (right) visits, split by region - after animals deviate from random-like sampling in session 1. **B** Same as in A, but for the whole of session 2. **C** Correlation of mean response to reward in a 1s period after a visit against reward rate (rewards/minute) at each option.

**Supp. Fig. 11.**
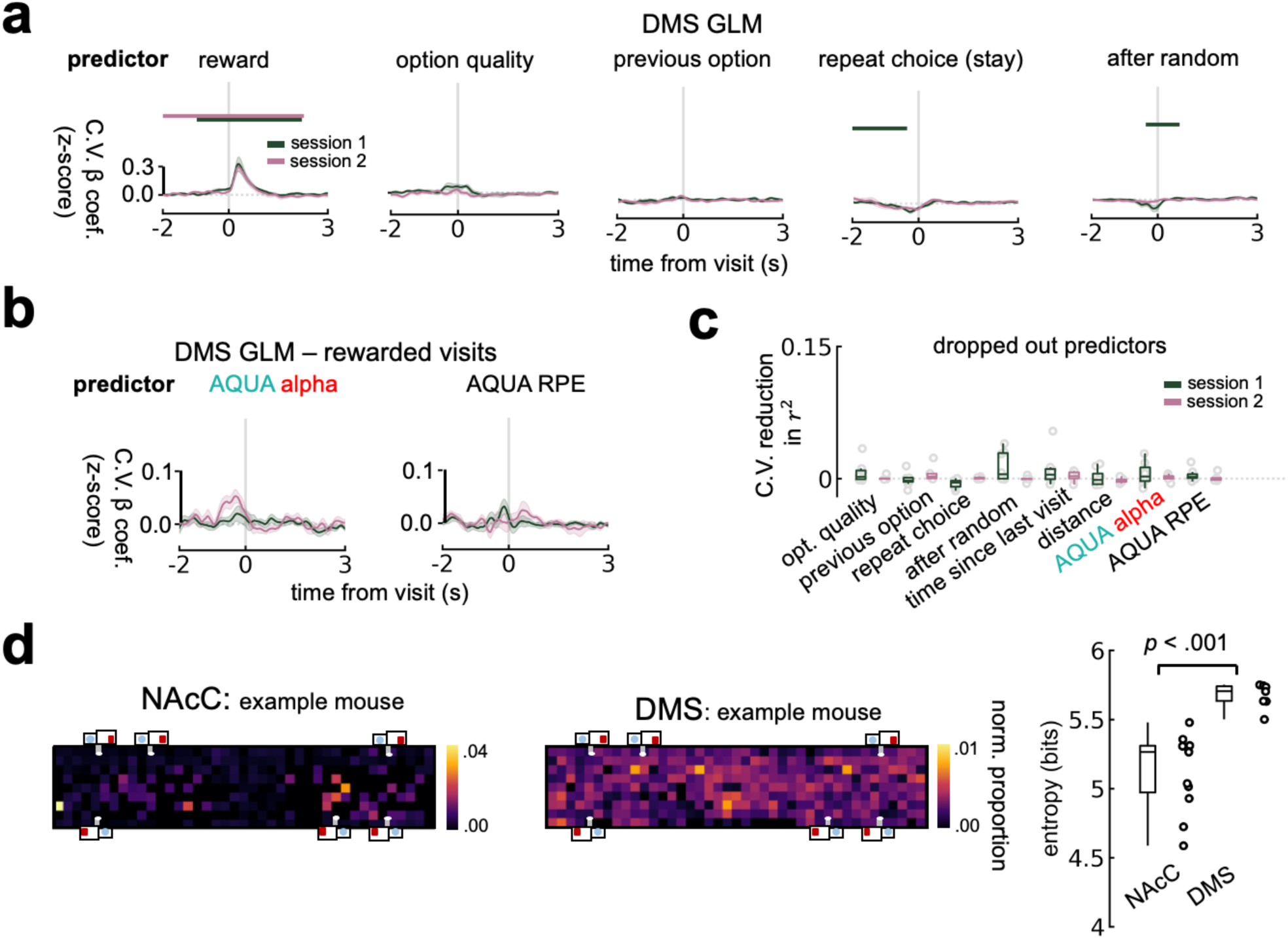
**A** GLM across rewarded and unrewarded visits for DMS signals. Bars indicate significant difference from zero (*p* < .01), Benjamini-Hochberg corrected for multiple comparisons. **B** AQUA alpha and RPE cross-validated beta coefficients from a GLM for DMS responses to rewarded visits. **C** 5-fold cross-validated reduction in GLM r2 with predictor drop out, predicting the mean of the peak response in the 1s period after reward delivery **D** Left: example proportions of identified transients in spatial bins, normalized by occupancy. All values in a given heatmap sum to 1. Right: entropy of transient distributions in space by region. Statistical test: two-tailed Mann Whitney U.

**Supp. Fig. 12.**
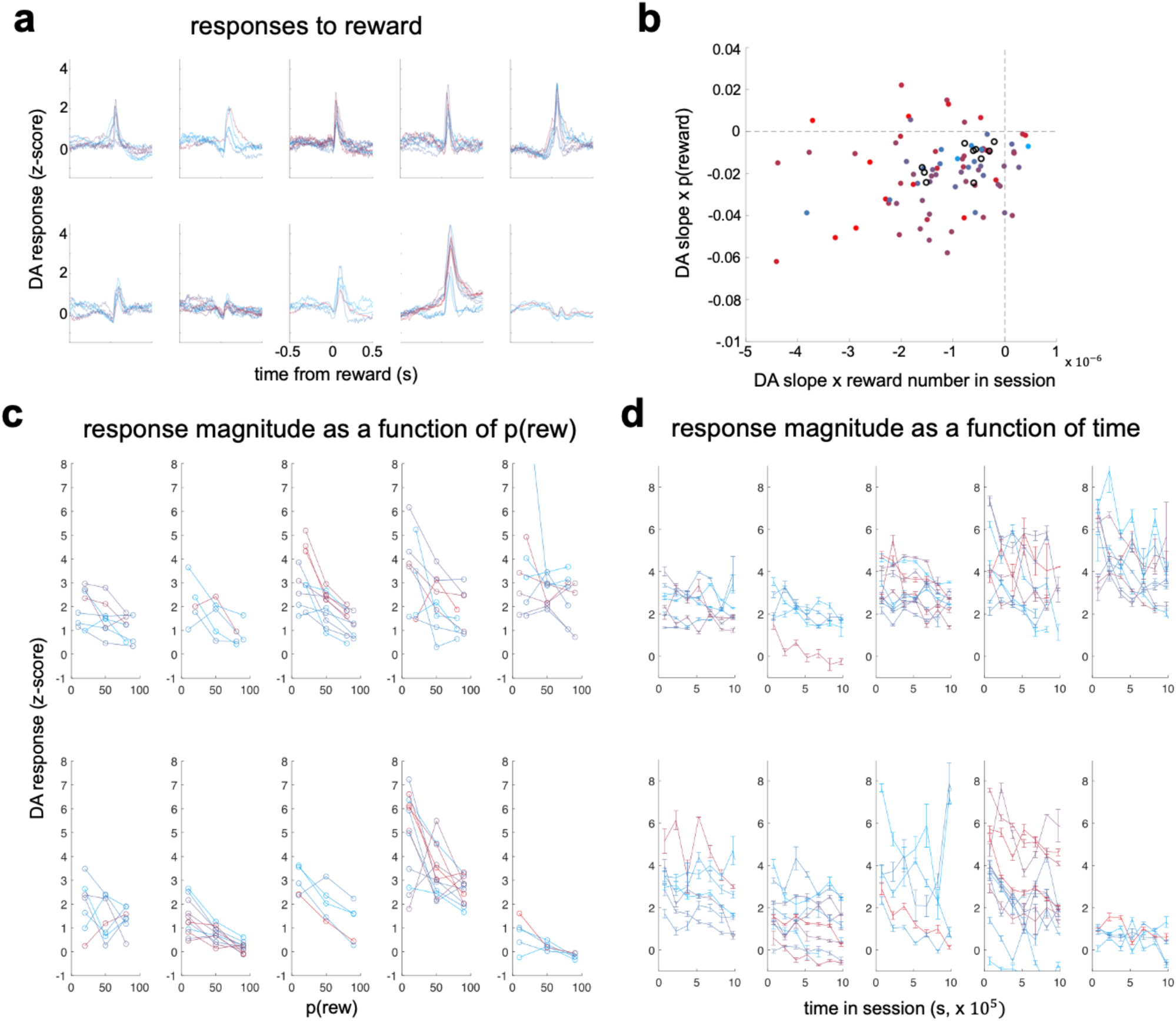
**A** Dopamine reward responses in a three-armed spatial bandit task [22] (one plot per mouse, colors indicate different sessions). **B** Slope of responses as a function of p(rew) against slope of responses as a function of reward number in session. Black circles are mean of slopes per mouse. All negative slopes are consistent with both or either a decaying learning rate or RPE. **C** Response magnitude per rat as a function of reward probability associated with each option. **D** Response magnitude per rat as a function of time in session. Note that slopes were fitted to lines in **C** and **D** produce **B**.

**Supp. Fig. 13.**
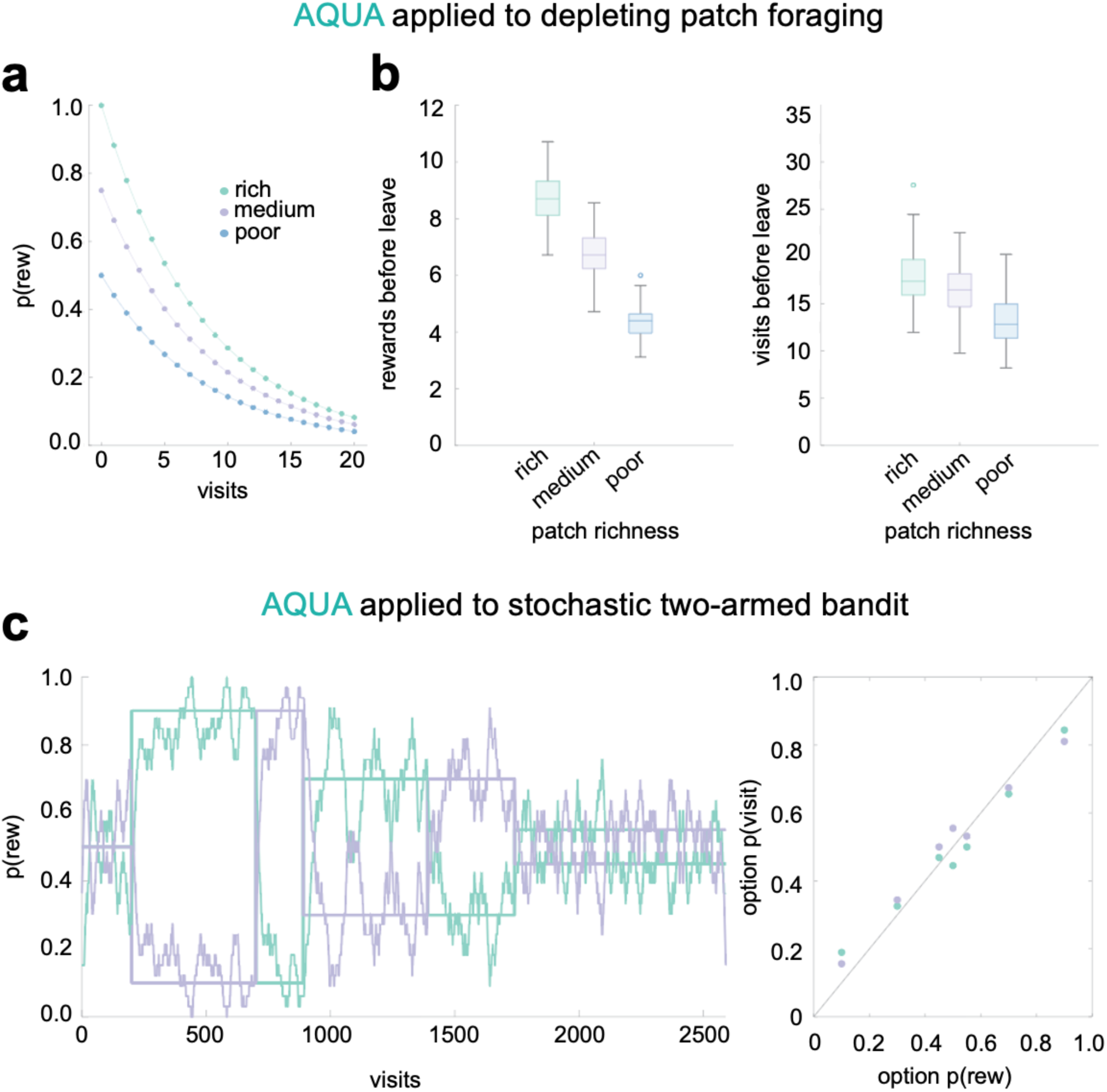
**A** Depleting patch foraging paradigm [9] where the probability of reward decays with repeated visits across three patches of differing quality. **B** Results from AQUA when run within this environment. Similar to MVT, AQUA harvests more rewards from rich patches than poor (left) and stays longer in rich patches (right). **C** AQUA behavior when run on stochastic two-armed bandit task [27] across visits (left), producing matching across options (right).

## Appendix 1

Here we discuss further insights from analysis of mouse treatment of space in FoMO. As discussed with reference to Fig. 3, the layout of options in FoMO meant that animals experienced 8 possible inter-spout distances, split across two ‘sides’ of the arena (Fig. 3a). Exploring the full range of possible pairwise transitions relative to the proportion of visits made at each option reveals that mice were sensitive to distance between options when choosing where to visit next (Fig. 3b), and this was also the case for animals that experienced config. 2 (Supp. Fig 6a). These mice were also less likely to make the direct transition from best to second best option, or vice versa - although with experience sensitivity to distance reduced (Supp. Fig. 6b). This treatment of distance appeared non-linear, as evidenced both by Fig. 3b/Supp. Fig. 6b and also by comparison of choice probabilities of the two matched options - 240s - in config. 1. Here sides of the arena contained the largest difference in average richness across options, yet strikingly mice chose them equally as often (Supp. Fig. 6c). This shows that animals were most sensitive to option quality *per se* over spatial distribution at intermediate spouts, reflecting their strong propensity to match even within the first session. This also reveals that in FoMO, mice treated evaluated each option separately rather than grouping them together in space as has been suggested as a mechanism for estimating average environment reward rate.

Thus far we have discussed the influence of objective distance in shaping animals’ decisions in FoMO. Critical to studies of foraging behavior and often incorporated into foraging theory is the importance of ‘travel’ when choosing to stay or select an alternative option. Here, where animals simultaneously have access to many possible options but are also totally unconstrained and free to execute variable trajectories, we had the unique opportunity to probe the relative importance of objective distance between options against the path length traveled by animals on their decisions of where to sample next. In practice travel times covering the different distances had highly overlapping distributions due to variability in path lengths, even when returning to the same spout without sampling any other (Supp. Fig. 6d). This variability was also seen across all objective distances between options (Supp. Fig. 6e). Therefore, whilst path length strongly correlated with travel time (Supp. Fig. 6f, left), objective distance between ports had a positive but weaker relationship to path length (Supp. Fig. 6f, middle) and travel time (Supp. Fig. 6f, right) as animals took variable and often indirect trajectories when traversing the FoMO arena.

As animals’ behavioral strategies evolved, path lengths relative to objective distance between options on average did decrease within the first session as animals moved away from pure exploration and quickly focused behavior on sampling the options available (Supp. Fig. 6g). However, on the second day despite animals rapidly settling into a matching strategy, path lengths increased (Supp. Fig. 6h). Taken together, these analyses suggest that animals were initially sensitive to distance between options but do not optimize path length with this degree of experience. This is in contrast to their proclivity to match when making decisions; in FoMO matching seems to emerge before path optimization.

## Acknowledgements

We thank L. Coddington and all members of the Dudman laboratory, Emily Dennis, and Alla Karpova for project feedback; J Talbot, S Sawtell, S Jager, and Janelia Experimental Technologies for technical assistance; the Janelia GENIE team for designing and supplying viral vectors; R. Demars, S. DiLisio, and S. Lindo for assistance with surgeries; M. Copeland, B. Foster, and M. Clarke for histology. This work was supported by the Howard Hughes Medical Institute, where J.T.D. and A.M.H. are Group Leaders at the Janelia Research campus.

## Author contributions

L.L.G. and J.T.D. conceived of the project and experiments, analyzed data, and wrote the manuscript with input from Y.G. and A.M.H. Y.G. designed and ran the optimality model with input from A.M.H. L.N. designed and implemented the moving gantry tracking camera system, including both custom hardware and software interface. J.T.D. and L.L.G. developed the AQUA computational model; J.T.D. ran the simulations. L.L.G. collected all experimental data.

## Data and code availability

Preliminary public versions of the code are available at: https://github.com/lauralgrima/hexaport_model

Additional code curation and hardware details will be available upon publication.

## References

1. Stephens DW, Krebs JR. Foraging Theory. Princeton University Press; 1986.

2. Charnov EL. Optimal foraging, the marginal value theorem. Theor Popul Biol. 1976;9: 129– 136.

3. Venkataraman VV, Kraft TS, Dominy NJ, Endicott KM. Hunter-gatherer residential mobility and the marginal value of rainforest patches. Proc Natl Acad Sci U S A. 2017;114: 3097– 3102.

4. Wajnberg E, Fauvergue X, Pons O. Patch leaving decision rules and the Marginal Value Theorem: an experimental analysis and a simulation model. Behav Ecol. 2000;11: 577– 586.

5. Turrin C, Fagan NA, Dal Monte O, Chang SWC. Social resource foraging is guided by the principles of the Marginal Value Theorem. Sci Rep. 2017;7: 11274.

6. Webb J, Steffan P, Hayden BY, Lee D, Kemere C, McGinley M. Foraging under uncertainty follows the marginal value theorem with Bayesian updating of environment representations. bioRxiv. 2024. doi:10.1101/2024.03.30.587253

7. Wajnberg, E., Bernhard, P., Hamelin, F. et al. Optimal patch time allocation for time-limited foragers. Behav Ecol Sociobiol. 2006;60: 1–10.

8. Vincent C, Frédéric G, Hamelin Frédéric M, Ludovic M. Taking fear back into the Marginal Value Theorem: the risk-MVT and optimal boldness. bioRxiv. 2024. p. 2023.10.31.564970. doi:10.1101/2023.10.31.564970

9. Lottem E, Banerjee D, Vertechi P, Sarra D, Lohuis MO, Mainen ZF. Activation of serotonin neurons promotes active persistence in a probabilistic foraging task. Nat Commun. 2018;9: 1000.

10. Kane GA, Bornstein AM, Shenhav A, Wilson RC, Daw ND, Cohen JD. Rats exhibit similar biases in foraging and intertemporal choice tasks. Elife. 2019;8. doi:10.7554/eLife.48429

11. Sylwestrak EL, Jo Y, Vesuna S, Wang X, Holcomb B, Tien RH, et al. Cell-type-specific population dynamics of diverse reward computations. Cell. 2022;185: 3568–3587.e27.

12. Pereira-Obilinovic U, Hou H, Svoboda K, Wang X-J. Brain mechanism of foraging: Reward-dependent synaptic plasticity versus neural integration of values. Proc Natl Acad Sci U S A. 2024;121: e2318521121.

13. Sutton RS, Barto AG. Reinforcement Learning: An Introduction. Cambridge, Mass.: MIT Press; 1998.

14. Grossman CD, Bari BA, Cohen JY. Serotonin neurons modulate learning rate through uncertainty. Curr Biol. 2022;32: 586–599.e7.

15. Costa VD, Dal Monte O, Lucas DR, Murray EA, Averbeck BB. Amygdala and ventral striatum make distinct contributions to reinforcement learning. Neuron. 2016;92: 505–517.

16. Schultz W, Dayan P, Montague PR. A neural substrate of prediction and reward. Science. 1997;275: 1593–1599.

17. Watabe-Uchida M, Eshel N, Uchida N. Neural circuitry of reward prediction error. Annu Rev Neurosci. 2017;40: 373–394.

18. Lak A, Stauffer WR, Schultz W. Dopamine neurons learn relative chosen value from probabilistic rewards. Elife. 2016;5: e18044.

19. Miller KJ, Botvinick MM, Brody CD. From predictive models to cognitive models: Separable behavioral processes underlying reward learning in the rat. bioRxiv. bioRxiv; 2018. doi:10.1101/461129

20. Baker S-A, Griffith T, Lepora NF. Degenerate boundaries for multiple-alternative decisions. Nat Commun. 2022;13: 5066.

21. Rosenberg M, Zhang T, Perona P, Meister M. Mice in a labyrinth: Rapid learning, sudden insight, and efficient exploration. 2021; 36.

22. Krausz TA, Comrie AE, Kahn AE, Frank LM, Daw ND, Berke JD. Dual credit assignment processes underlie dopamine signals in a complex spatial environment. Neuron. 2023;111: 3465–3478.e7.

23. Li Y, Dudman JT. Mice infer probabilistic models for timing. Proc Natl Acad Sci U S A. 2013;110: 17154–17159.

24. Soares S, Atallah BV, Paton JJ. Midbrain dopamine neurons control judgment of time. Science. 2016;354: 1273–1277.

25. Herrnstein RJ. Relative and absolute strength of response as a function of frequency of reinforcement. J Exp Anal Behav. 1961;4: 267–272.

26. Tsutsui K-I, Grabenhorst F, Kobayashi S, Schultz W. A dynamic code for economic object valuation in prefrontal cortex neurons. Nat Commun. 2016;7: 12554.

27. Sugrue LP, Corrado GS, Newsome WT. Matching behavior and the representation of value in the parietal cortex. Science. 2004;304: 1782–1787.

28. Rajagopalan AE, Darshan R, Fitzgerald JE, Turner GC. Expectation-based learning rules underlie dynamic foraging in Drosophila. bioRxiv. 2022. p. 2022.05.24.493252. doi:10.1101/2022.05.24.493252

29. Baum WM. On two types of deviation from the matching law: bias and undermatching. J Exp Anal Behav. 1974;22: 231–242.

30. Herrnstein R. The matching law: Papers in psychology and economics. Rachlin H, Laibson DI, editors. New York, NY, US: Russell Sage Foundation; Cambridge, MA, US; 1997.

31. Vulkan N. An economist’s perspective on probability matching. J Econ Surv. 2000;14: 101– 118.

32. Corrado G, Doya K. Understanding neural coding through the model-based analysis of decision making. J Neurosci. 2007;27: 8178–8180.

33. Barack DL. What is foraging? Biol Philos. 2024;39: 3.

34. Sakai Y, Fukai T. The actor-critic learning is behind the matching law: matching versus optimal behaviors. Neural Comput. 2008;20: 227–251.

35. Loewenstein Y, Seung HS. Operant matching is a generic outcome of synaptic plasticity based on the covariance between reward and neural activity. Proc Natl Acad Sci U S A. 2006;103: 15224–15229.

36. Kingma DP, Ba J. Adam: A Method for Stochastic Optimization. arXiv [cs.LG]. 2014. Available: http://arxiv.org/abs/1412.6980

37. Mnih V, Kavukcuoglu K, Silver D, Graves A, Antonoglou I, Wierstra D, et al. Playing Atari with Deep Reinforcement Learning. arXiv [cs.LG]. 2013. Available: http://arxiv.org/abs/1312.5602

38. Coddington LT, Lindo SE, Dudman JT. Mesolimbic dopamine adapts the rate of learning from action. Nature. 2023;614: 294–302.

39. Watkins CJCH, Dayan P. Q-learning. Mach Learn. 1992;8: 279–292.

40. Engelhard B, Finkelstein J, Cox J, Fleming W, Jang HJ, Ornelas S, et al. Specialized coding of sensory, motor and cognitive variables in VTA dopamine neurons. Nature. 2019;570: 509–513.

41. Syed ECJ, Grima LL, Magill PJ, Bogacz R, Brown P, Walton ME. Action initiation shapes mesolimbic dopamine encoding of future rewards. Nat Neurosci. 2016;19: 34–36.

42. Berke JD. What does dopamine mean? Nat Neurosci. 2018;21: 787–793.

43. Burwell SCV, Yan H, Lim SSX, Shields BC, Tadross MR. Natural phasic inhibition of dopamine neurons signals cognitive rigidity. bioRxiv. 2024. doi:10.1101/2024.05.09.593320

44. Bukwich M, Campbell MG, Zoltowski D, Kingsbury L, Tomov MS, Stern J, et al. Competitive integration of time and reward explains value-sensitive foraging decisions and frontal cortex ramping dynamics. bioRxiv. 2023. doi:10.1101/2023.09.05.556267

45. McNamara JM, Houston AI. Optimal foraging and learning. J Theor Biol. 1985;117: 231– 249.

46. Kilpatrick ZP, Davidson JD, Hady AE. Normative theory of patch foraging decisions. bioRxiv. bioRxiv; 2020. doi:10.1101/2020.04.22.055558

47. Constantino SM, Daw ND. Learning the opportunity cost of time in a patch-foraging task. Cogn Affect Behav Neurosci. 2015;15: 837–853.

48. Shahidi N, Franch M, Parajuli A, Schrater P, Wright A, Pitkow X, et al. Population coding of strategic variables during foraging in freely moving macaques. Nat Neurosci. 2024;27: 772– 781.

49. Wittmann MK, Kolling N, Akaishi R, Chau BKH, Brown JW, Nelissen N, et al. Predictive decision making driven by multiple time-linked reward representations in the anterior cingulate cortex. Nat Commun. 2016;7: 12327.

50. Kolling N, Akam T. (Reinforcement?) Learning to forage optimally. Curr Opin Neurobiol. 2017;46: 162–169.

51. Crowcroft P. Mice all over. Chicago Zoological Society; 1973.

52. Sousa M, Bujalski P, Cruz BF, Louie K, McNamee D, Paton JJ. Dopamine neurons encode a multidimensional probabilistic map of future reward. bioRxiv. 2023. doi:10.1101/2023.11.12.566727

53. Balsam PD, Gallistel CR. Temporal maps and informativeness in associative learning. Trends Neurosci. 2009;32: 73–78.

54. Loewinger G, Cui E, Lovinger D, Pereira F. A statistical framework for analysis of trial-level temporal dynamics in fiber photometry experiments. 2024. doi:10.7554/elife.95802

55. Akam T, Lustig A, Rowland JM, Kapanaiah SK, Esteve-Agraz J, Panniello M, et al. Open-source, Python-based, hardware and software for controlling behavioural neuroscience experiments. Elife. 2022;11: e67846.

56. Akam T, Walton ME. pyPhotometry: Open source Python based hardware and software for fiber photometry data acquisition. Sci Rep. 2019;9: 3521.

57. Paxinos G, Franklin KBJ. Paxinos Franklin’s Mouse Brain Stereotaxic Coordinates (Academic). 2019.

